# Metabolic-State-Dependent Magnetic Field Effects on Endogenous Flavin Autofluorescence in Escherichia coli

**DOI:** 10.64898/2026.10.23.761314

**Authors:** Noboru Ikeya, Mio Omi, Jonathan R. Woodward

## Abstract

We demonstrate that endogenous autofluorescent molecules in Escherichia coli exhibit fluorescence changes exceeding 10% in response to a 25 mT magnetic field under specific physiological and excitation conditions. Fluorescence spectral analysis strongly supports a flavin origin of the magnetically sensitive signal, with a slight blue shift relative to free flavin suggesting a protein-associated population. The analysis further suggests that this magnetic response originates from a flavin-related component contributing only part of the total cellular autofluorescence. This implies that the magnetic field effect exceeding 10% observed in total cellular autofluorescence represents an ensemble-averaged response, and that the intrinsic magnetic field effect within the magnetically sensitive flavin population may be substantially larger. Notably, the magnetic field effect is strongly suppressed by carbon-source supplementation. These findings indicate that the manifestation of the magnetic field effect is associated with the cellular physiological or metabolic state. The field-induced negative fluorescence response and the saturation of the magnetic-field dependence support the radical-pair mechanism and are consistent with a triplet-born radical-pair reaction. This work provides a framework for identifying endogenous biomolecular candidates exhibiting large intracellular magnetic field effects and suggests a new route toward label-free readout of cellular metabolic states based on flavin magnetosensitivity.

## Introduction

Biological effects of weak magnetic fields have been reported across diverse phenomena, including magnetoreception, circadian rhythms, cell growth, redox responses and neuronal function^1^. A central challenge in this field is to connect magnetically sensitive molecular processes to cellular responses and physiological function. The radical-pair mechanism (RPM) provides a leading molecular-level basis for explaining how weak magnetic fields can influence biochemical reactions^1,2^. In this mechanism, magnetic-field-dependent singlet– triplet interconversion in radical pairs generated by photochemical or redox reactions can alter reaction yields or reaction branching, providing a route by which weak magnetic fields, despite involving interaction energies far smaller than thermal energy, can influence biochemical outcomes^1–3^. A substantial gap nevertheless remains between molecular-level spin chemistry and cellular-level physiological responses.

Cryptochrome is the most extensively studied example of flavin-dependent magnetic-field sensitivity in biology^2,4–7^. Spectroscopic and biochemical studies have defined its FAD photochemistry, radical-pair formation, spin-dependent reaction pathways, and magnetic-field sensitivity^8–11^. Cryptochrome has also been implicated in biological functions including ROS signaling^12^, neuronal excitability^13^, circadian regulation^14^, and magnetosensory behavior^15^. However, cryptochrome represents only one part of a broader cellular flavin landscape: flavins are widely present in cells as free molecules and as bound molecules in diverse flavoproteins^16,17^, and RPM-based magnetic-field effects (MFEs) have also been observed in non-cryptochrome flavins and model reaction systems^18^. An important question is therefore not simply whether flavin-related molecules can exhibit MFEs in cells, but which of these potentially diverse flavin-based radical-pair reactions, in which subcellular locations, cellular states, and molecular environments, acquire functional significance and manifest as observable magnetic-field sensitivity.

A first step toward addressing this question is to establish how endogenous flavin-related magnetic-field sensitivity depends on cellular state. A spectroscopic approach that probes endogenous flavin states together with their magnetic-field responses provides a means of examining this relationship in living cells. Oxidized flavins emit green autofluorescence under blue-light excitation, and autofluorescence arising from FAD, FMN, riboflavin, and flavoprotein cofactors therefore provides an endogenous, label-free signal for probing intracellular flavin states^19,20^. This signal can reflect not only flavin abundance but also cell-state-dependent information such as redox state, protein-binding state, local molecular environment, and intracellular localization^21,22^. Furthermore, by analyzing the MFE on this autofluorescence, it is possible to test whether endogenous flavin signals contain spin-dependent reaction components involving radical-pair formation^23^. Thus, endogenous flavin autofluorescence MFE measurements provide a powerful approach for detecting and evaluating latent flavin-based radical-pair reactions in living cells by combining sensitivity to cellular state with selectivity for spin-dependent reactions.

However, connecting the magnetic sensitivity of flavin molecules to cellular function through endogenous flavin autofluorescence MFE measurements remains technically challenging in living cells. We previously observed autofluorescence MFEs in living HeLa cells using non-confocal wide-field imaging^23^. By integrating fluorescence over a broad observation volume, including regions outside the focal plane, this approach reduced intensity fluctuations associated with intracellular movement of fluorescent structures, enabling sensitive detection of small magnetic-field responses in individual cells. However, the limited illuminated area of that setup constrained simultaneous measurements across many cells and systematic exploration of experimental conditions. In contrast, another group used spinning-disk confocal microscopy to statistically evaluate autofluorescence MFEs across multiple cells, but detected no significant MFE under their experimental conditions^24^. Although this approach allows scanning over a wide field of view, that study identified weak autofluorescence signals, intensity fluctuations associated with intracellular movement, and cell-to-cell variability as challenges to detection. Furthermore, because flavin-based MFEs can depend strongly on excitation intensity and duration^25,26^, spatial uniformity of and intensity of excitation must also be considered and measured when comparing and quantifying magnetic-field responses across cells. Therefore, ideally a complementary experimental system that enables simultaneous measurements across many cells under uniform excitation while maintaining high detection sensitivity is desirable.

*Escherichia coli* (*E. coli*) is a suitable model system for such measurements. Because *E. coli* cells are small, magnetic-field responses can be measured simultaneously across many cells at a sufficiently high and relatively uniform excitation intensity, without enlarging the illuminated area or requiring a corresponding increase in laser power. This enables assessment of population-level magnetic-field responses and systematic comparisons across experimental conditions while maintaining high detection sensitivity in each measurement. Furthermore, *E. coli* also exhibits green autofluorescence, to which flavin-related molecules including FAD, FMN, riboflavin, and flavoprotein cofactors are expected to make a substantial contribution^27,28^. In addition, *E. coli* is easy to culture and its physiological state can be readily controlled, facilitating reproducible investigation of how endogenous flavin autofluorescence MFEs depend on cellular state.

In this study, we demonstrate a clear endogenous autofluorescence MFE in living *E. coli* populations. Wavelength-resolved MFE analysis suggests that this response originates from a flavin-related component contributing only part of the total cellular autofluorescence. We further show that the manifestation of this MFE is associated with cellular physiological or metabolic state, as evidenced by its suppression upon glucose addition and delayed recovery after glucose removal. Metabolism connects environmental inputs to cellular functions through biochemical transformations and energy conversion, thereby providing an intermediate layer between molecular-level reaction processes and cell-level physiological outputs. Thus, by manipulating nutrient conditions, this study presents a new experimental framework for revealing endogenous flavin-related MFEs and examining their dependence on cellular state.

## Results

### Magnetic Field Effects on Endogenous Autofluorescence in *E. coli*

We first examined whether a clear magnetic field effect could be observed at the population level in the endogenous autofluorescence of *E. coli*. A simple, standard sample-preparation protocol was used: cells grown in LB broth were washed and resuspended in PBS (**Fig. 1A**). Fluorescence was measured using a custom-built microscope (**Fig. S1**).

**Figure 1.**
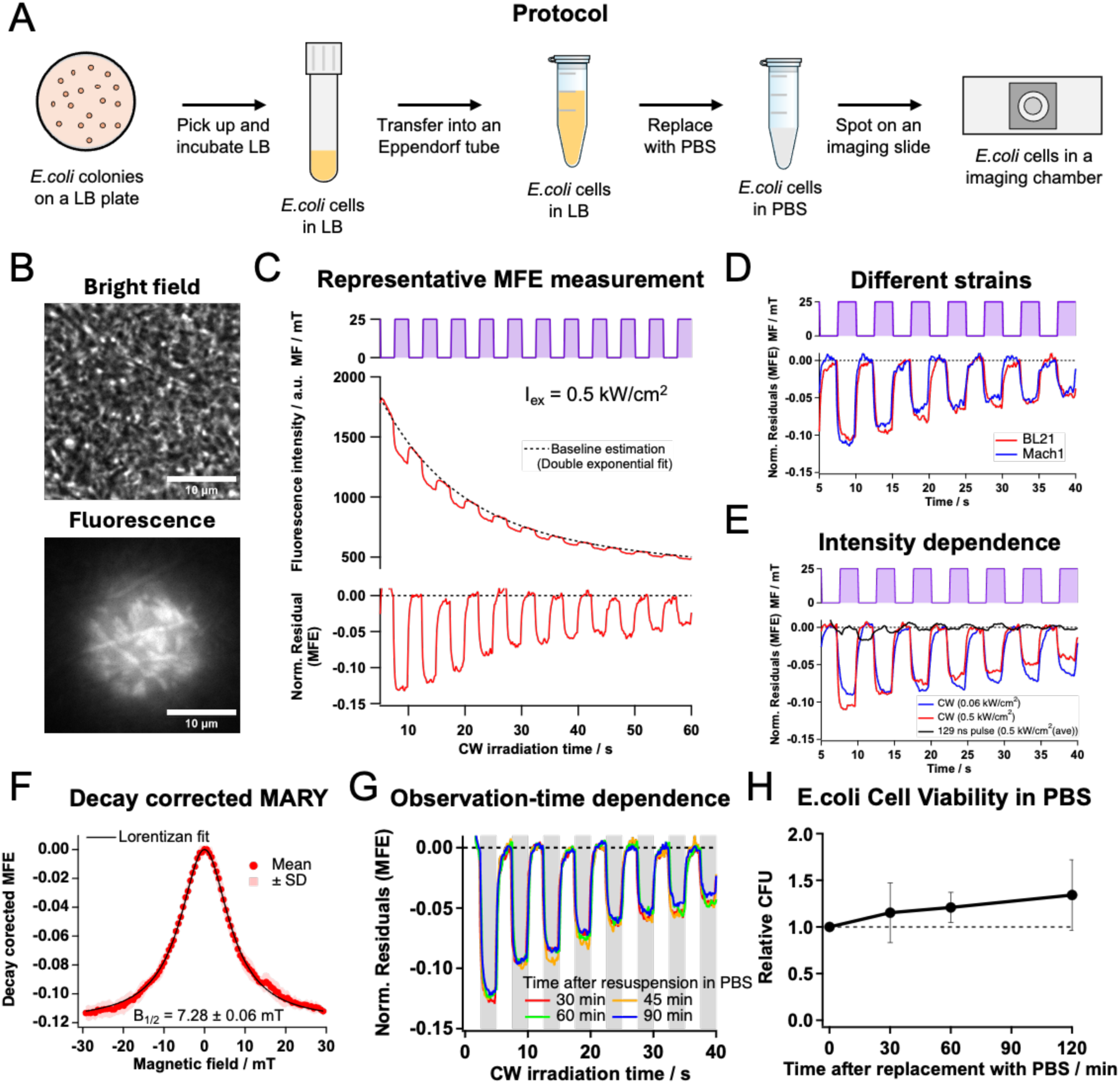
Magnetic Field Effects (MFE) on Endogenous Autofluorescence in *E. coli*. (A) Schematic of the sample preparation protocol. *E. coli* cells grown in LB broth were washed and resuspended in phosphate-buffered saline (PBS). The suspension was mounted between two glass coverslips using an imaging spacer (see the Method section). (B) Representative bright-field (top) and autofluorescence (bottom) images of an *E. coli* population in PBS (OD₆₀₀ = 14.3). Scale bars, 10 μm.(C) Representative measurement of the MFE on autofluorescence under continuous-wave (CW) excitation at an excitation intensity of (Iex = 0.5 kW/cm^2^). Top: applied magnetic field modulated between 0 and 25 mT. Middle: mean fluorescence intensity (F(t)) within the region of interest (ROI) (red line) and fitted bi-exponential baseline (black dotted line). Bottom: normalized fluorescence residual, ((F(t)-Ffit(t))/Ffit(t)), corresponding to the MFE. See Supporting Information for details of baseline estimation. (D) Autofluorescence MFEs in two *E.coli* strains (BL21 and Mach1). (E) Excitation intensity dependence of representative autofluorescence MFEs on BL 21. Cells were excited with CW light at 0.06 or 0.5 kW/cm² or with pulsed light (129 ns pulse width, 32 kHz repetition rate) at a time-averaged intensity of 0.5 kW/cm². (F) Decay-corrected magnetically affected reaction yield (MARY) curve for autofluorescence from *E. coli* BL21 cells, measured using 0.05-Hz triangular-wave modulation between −29.25 and +29.25 mT at an excitation intensity of 0.06 kW/cm². Data show the mean ± SD of six consecutive 10-s sweeps acquired 15–75 s after excitation onset, following correction for fluorescence baseline decay and MFE amplitude decay. A Lorentzian fit to the mean curve is shown. See Fig. S4 for the analysis procedure. (G) Time course of the autofluorescence MFE in *E. coli* cells suspended in PBS. (H)Viable cell counts of *E. coli* (BL21) after resuspension in PBS. CFU mL⁻¹ values were normalized to time 0 within each biological replicate. Data are mean ± SD (n = 3 biological replicates).

**Fig. 1B** shows representative bright-field and autofluorescence images of an *E. coli* population used for MFE measurements (approximately 300 cells; see SI for estimated cell numbers within the excitation region). During irradiation, the mean fluorescence intensity within the region of interest (ROI) showed an overall decay (**Fig. 1C**). Superimposed on this decay, on/off modulation of a 25 mT magnetic field produced clear fluorescence decreases when the field was on and recovery toward the decaying baseline when it was off. To quantify the relative fluorescence change (MFE), we estimated the baseline by least-squares fitting with a bi-exponential function, then subtracted it from the measured intensity and normalized the residuals to the baseline (see SI for details). Under these excitation conditions, magnetic field application reduced fluorescence by more than 10%. Similar MFEs were observed in both BL21 and Mach1 strains (**Fig. 1D**), indicating that the phenomenon is not restricted to a single strain.

We next examined how the MFE depended on excitation conditions (**Fig. 1E**). Higher-intensity continuous-wave (CW) excitation produced a larger MFE amplitude but more pronounced decay during irradiation than lower-intensity CW excitation; the decay was even more pronounced under pulsed excitation. To characterize the magnetic-field dependence, we performed magnetically affected reaction yield (MARY) measurements using triangular-wave modulation between −29.25 and +29.25 mT, with each sweep lasting 10 s, at a reduced excitation intensity of 0.06 kW/cm² (compared with 0.5 kW/cm² for the measurements above). We developed a MARY analysis procedure to correct for the irradiation-induced decay of the MFE amplitude. Following fluorescence baseline correction, the response was normalized using a decay envelope estimated from the responses at the magnetic-field extrema. Six consecutive sweeps acquired 15–75 s after excitation onset were then aligned on a common magnetic-field axis and averaged to obtain the decay-corrected MARY curve (**Fig. 1F**; see **Fig. S4** and SI for analysis details). The MFE magnitude increased with magnetic-field strength and approached saturation. A Lorentzian fit to the corrected mean curve yielded a half-field value of B₁/₂ = 7.28 ± 0.06 mT. These dependences on excitation intensity and magnetic-field strength are consistent with a radical-pair origin of the fluorescence MFE.

The MFE measurements were performed within approximately 1.5 h after suspension in PBS, and the MFE was repeatedly observed throughout this period (**Fig.1G**). To determine whether *E. coli* remained viable after suspension in PBS under the protocol shown in **Fig.1A**, culturability was evaluated. As shown in **Fig.1H**, little change in viable cell count was observed even after 2 h in PBS. Consistent with this observation, previous studies have likewise shown that the viable cell counts of *E. coli* K-12 remain nearly constant for at least the first 24 h in PBS, regardless of incubation temperature (5 °C or 30 °C)^29^.

Taken together, these results show that endogenous autofluorescence from viable *E. coli* populations exhibits a robust magnetic field effect in PBS, with a fluorescence decrease exceeding 10% at 25 mT under 450 nm excitation. The saturating field dependence is characteristic of a radical-pair mechanism, and the negative sign of the response is that expected for a triplet-born radical pair.

### 2. Spectral Characterization of the Magnetic-Field-Sensitive Autofluorescence in *E. coli*

Cellular autofluorescence is a composite signal arising from multiple endogenous fluorophores and their microenvironment-dependent emission properties^30^. Therefore, to characterize the molecular components contributing to the MFE observed in the previous section, we analyzed the total fluorescence spectrum, the illumination-induced difference fluorescence spectrum, and the magnetic-field-induced difference fluorescence spectrum using *E. coli* cells prepared under the same culture conditions.

The total autofluorescence spectra, corrected by subtraction of an independently measured PBS background spectrum, progressively decreased in intensity during continuous 450 nm excitation (**Fig. 2A**). After normalization to their respective peak intensities, the spectra at different irradiation times closely overlapped (**Fig. 2B**), indicating that the overall spectral profile was largely preserved as the fluorescence intensity decreased. The illumination-induced difference fluorescence spectrum, F(λ,t)-F(λ,0), was calculated to extract spectral changes occurring during irradiation. Because time-invariant contributions are canceled by subtraction, contributions from scattered light, PBS-derived background, and fluorescence components that remain nearly constant during irradiation are strongly suppressed. This enables fluorescence changes induced by 450 nm excitation to be evaluated more directly. A broad negative difference signal developed in the main autofluorescence region and increased in absolute magnitude with irradiation time (**Fig. 2C**). When normalized to the magnitude of their respective negative peaks, the difference spectra obtained at different irradiation times showed highly similar spectral profiles (**Fig. 2D**). This indicates that the spectral profile of the fluorescence lost during irradiation remained approximately constant over time.

**Figure 2.**
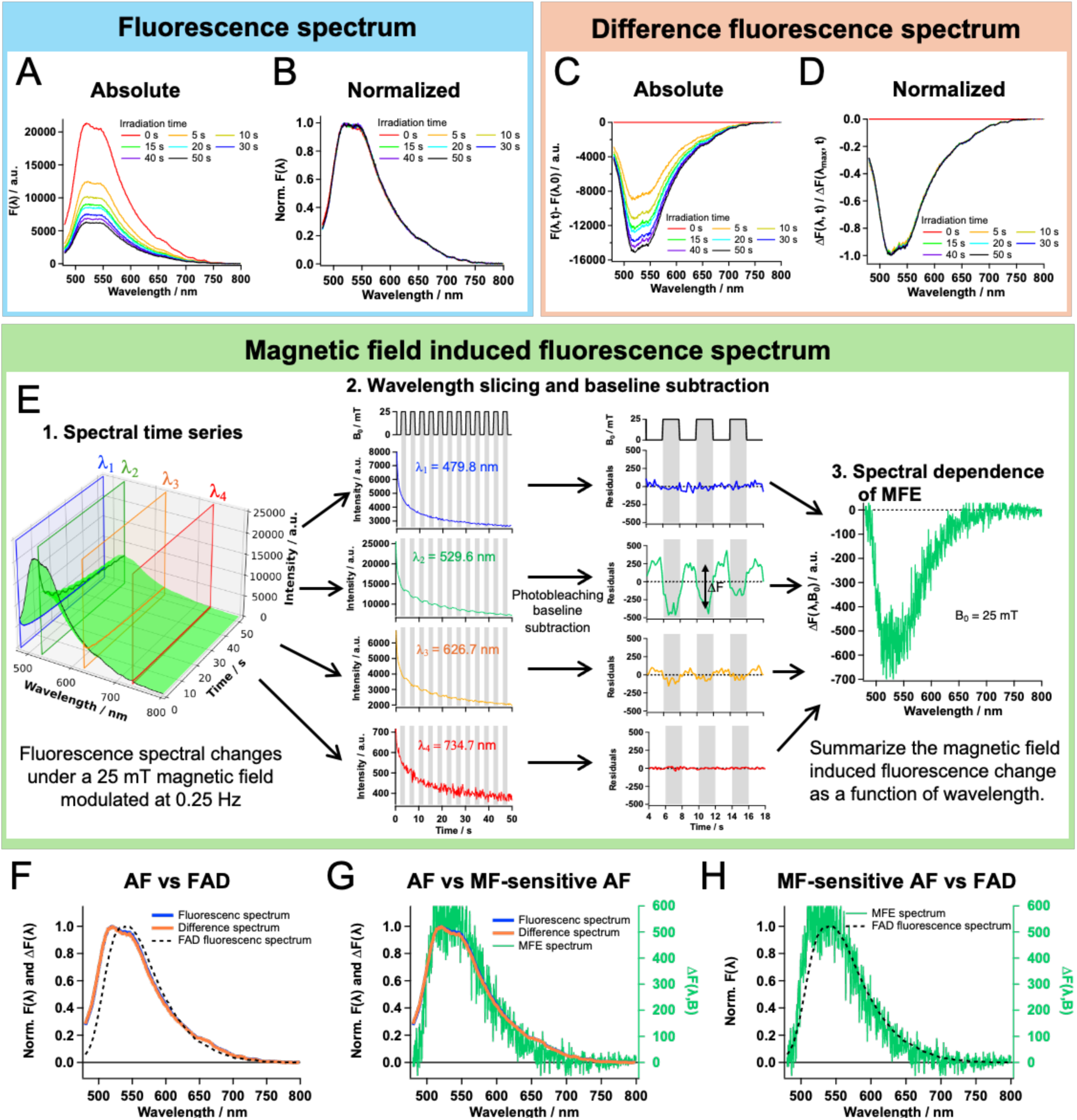
Spectral Characterization of the Magnetic-Field-Sensitive Autofluorescence in *E. coli* BL21. (A) Total autofluorescence spectra. Spectra were recorded at irradiation times of 0, 5, 10, 15, 20, 30, 40, and 50 s under continuous 450 nm excitation (Iex = 0.5 kW/cm^2^). (B) Normalized total autofluorescence spectra from (A). Each spectrum was normalized to its peak intensity. (C) Difference fluorescence spectra from (A). The illumination-induced difference spectra were calculated as (F(λ,t)-F(λ,0)), where λ and t denote wavelength and irradiation time, respectively. (D) Normalized difference fluorescence spectra from (C). Each spectrum was normalized to the magnitude of its negative peak. (E) Magnetic-field-induced difference fluorescence spectrum (wavelength-resolved MFE spectrum). The spectrum was obtained under on/off modulation of a 25 mT magnetic field by applying baseline subtraction and residual analysis to the fluorescence decay trace at each wavelength, as in Fig. 1C. (F) Comparison of the total autofluorescence spectrum and the difference fluorescence spectrum with the fluorescence spectrum of FAD in PBS buffer. (G) Comparison of the total autofluorescence spectrum, the difference fluorescence spectrum, and the wavelength-resolved MFE spectrum. (H) Comparison of the wavelength-resolved MFE spectrum with the fluorescence spectrum of FAD in PBS buffer.

To determine the wavelength dependence of the magnetic-field-induced fluorescence change, fluorescence spectra were recorded under on/off modulation of a 25 mT magnetic field. The fluorescence traces contained both magnetic-field-independent photobleaching and magnetic-field-dependent responses. Because the MFE itself decreased during irradiation, the analysis was restricted to a short measurement window. Within this window, residual analysis, as in **Fig. 1C**, was performed at each wavelength point to subtract the photobleaching baseline and determine the wavelength-resolved MFE. As shown in **Fig. 2E**, this analysis was performed over the 480–800 nm range at wavelength intervals of approximately 0.4 nm. The residual signal at each wavelength was then used to construct the magnetic-field-induced difference fluorescence spectrum, hereafter referred to as the wavelength-resolved MFE spectrum. Notably, no clear MFE was detected in the short-wavelength region, despite the presence of measurable fluorescence and illumination-induced fluorescence decay in this region. This indicates that the MFE was not simply proportional to, or a general consequence of, the illumination-induced fluorescence decay. Instead, the wavelength dependence indicates that the MFE is associated with only a subset of the autofluorescent spectral components and is less consistent with a wavelength-independent background or a simple global intensity artifact. Both spectra also exhibited a small excess signal in the long-wavelength region compared with free FAD.

We next compared the total fluorescence spectrum, the illumination-induced difference fluorescence spectrum, the wavelength-resolved MFE spectrum, and the fluorescence spectrum of FAD in solution (**Fig.2F–H**). As shown in **Fig.2F**, the total fluorescence spectrum and the illumination-induced difference fluorescence spectrum were clearly shifted toward shorter wavelengths relative to free FAD. Previous studies have shown that protein-bound or microenvironmentally constrained flavins can exhibit fluorescence spectra shifted toward shorter wavelengths compared with free flavins in solution^31,32^. In addition, green autofluorescence in *E. coli* has been reported to arise mainly from flavin-related molecules, including FAD, FMN, and riboflavin^27,28^, and *E. coli* contains numerous flavoproteins with FAD or FMN cofactors^33^. Therefore, the blue-shifted profiles of the total fluorescence and illumination-induced difference spectra are consistent with contributions from flavins in protein-bound or otherwise microenvironmentally constrained states.

In contrast, the wavelength resolved MFE spectrum was not identical to the total fluorescence spectrum, the illumination-induced difference fluorescence spectrum, or the spectrum of free FAD in solution (**Fig.2G, H**). As shown in **Fig.2G**, the total fluorescence spectrum and the illumination-induced difference fluorescence spectrum showed clear signals in the short-wavelength region, whereas the contribution of this region was minimal in the wavelength-resolved MFE spectrum, similar to free FAD. However, as shown in **Fig.2H**, the wavelength-resolved MFE spectrum was not identical to free FAD and was slightly shifted toward shorter wavelengths. These results suggest that the MFE does not arise uniformly from all endogenous autofluorescent molecules in *E.coli*, but is instead associated with a specific flavin-related component. Furthermore, its spectral profile is consistent with contributions from flavin species that weakly interact with, or are loosely associated with, intracellular macromolecular environments.

Together, these spectral analyses indicate that the magnetic-field-sensitive autofluorescence in *E. coli* is flavin-like, but spectrally distinct from both the total autofluorescence and free FAD in solution. Under the present 450 nm excitation condition, the most plausible molecular candidates are oxidized flavin-related species, including FAD, FMN and riboflavin, with FAD and FMN potentially bound within proteins or interacting with protein surfaces. These results support the idea that the observed MFE is associated with a specific subset of intracellular flavin-related components, rather than the entire autofluorescent molecular pool.

### 3. Glucose Dependence of the Magnetic Field Effect on Autofluorescence in *E. coli*

While PBS provides a chemically simple environment that minimizes nutrient-derived confounding factors, it also places *E. coli* under nutrient-deprived conditions. We therefore next examined the autofluorescence MFE under nutrient-containing conditions. Notably, the MFE was markedly suppressed under glucose-containing conditions.

We first examined the MFE in M9 minimal medium supplemented with glucose, a chemically defined medium that provides glucose and defined nutrients supporting bacterial metabolism. Under this condition, the MFE was markedly reduced (**Fig. 3A**). To determine whether this suppression was specific to M9 medium or associated with glucose supplementation, we next measured the MFE in PBS supplemented with glucose. The MFE was also markedly reduced in PBS + glucose, whereas a clear negative MFE was observed in PBS alone under the same excitation and magnetic-field conditions (**Fig. 3A**). At the lower excitation intensity (0.06 kW/cm²), a small residual MFE remained detectable in PBS + glucose, whereas at the higher excitation intensity (0.5 kW/cm²), the MFE was nearly abolished. We therefore quantified the MFE amplitude at 0.06 kW/cm², where the MFE showed less decay during irradiation, and found a significant suppression of the autofluorescence MFE in PBS + glucose compared with PBS alone (**Fig. 3B**). These results indicate that MFE suppression is consistently associated with glucose-containing conditions in both M9 and PBS backgrounds, despite their distinct buffer and medium compositions.

**Figure 3.**
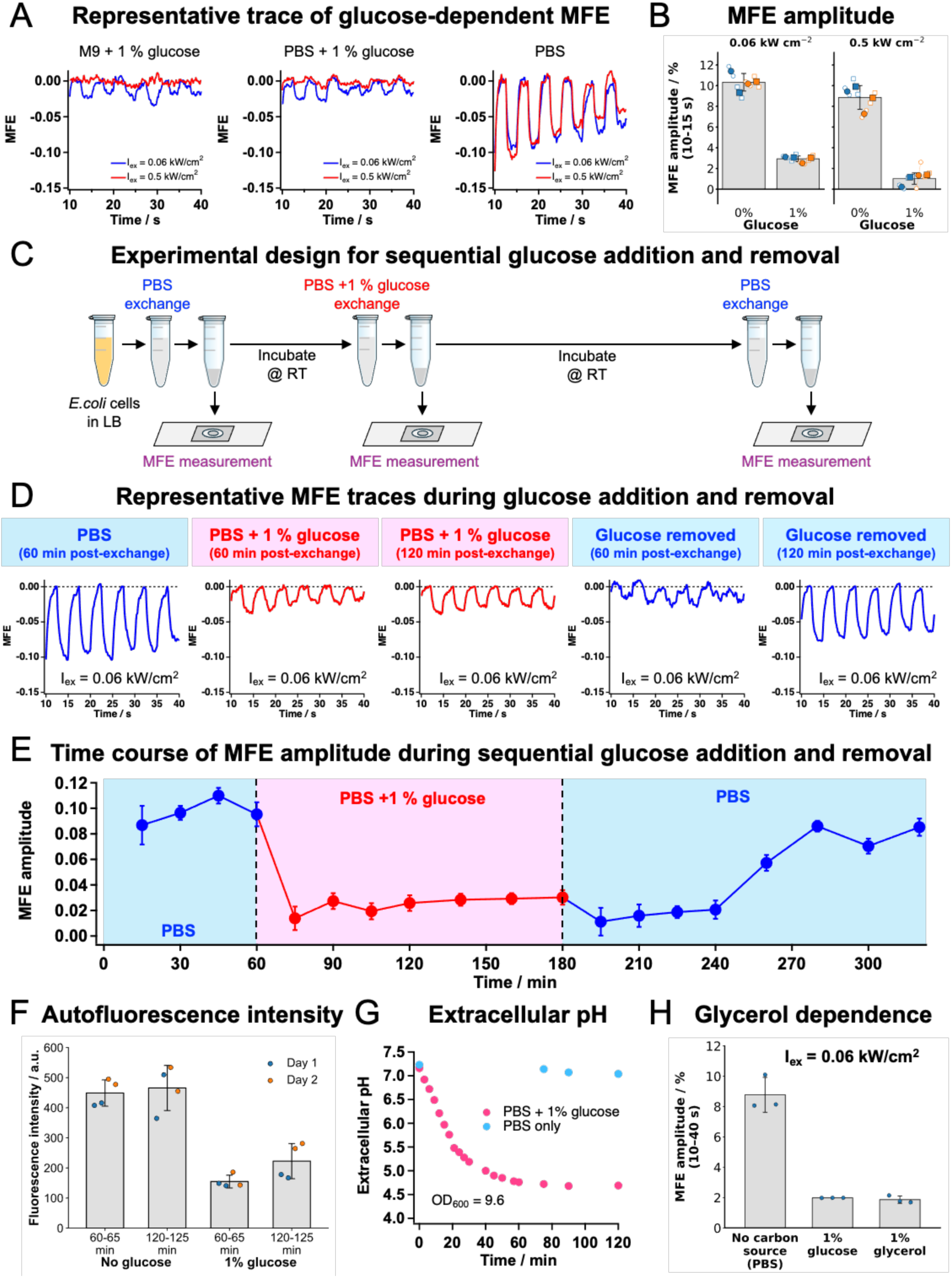
Glucose Dependence of the Magnetic Field Effect on Autofluorescence in *E. coli* (BL21). (A) Representative traces of the glucose-dependent autofluorescence MFE of *E.coli* population (OD600 = 10.0). Traces were recorded in PBS, PBS supplemented with 1% (w/v) glucose, and M9 minimal medium supplemented with 1% glucose at excitation intensities of 0.06 or 0.5 kW cm⁻², as indicated. (B) Comparison of the MFE amplitude in PBS with and without glucose supplementation. (C) Experimental design for sequential glucose addition and removal. The medium was exchanged sequentially from LB to PBS, then to PBS supplemented with 1% glucose, and finally back to PBS, with incubation at room temperature between exchanges. After each exchange, an aliquot was taken for slide preparation, while the remaining suspension continued through the sequence in the tube. At each stage, incubation in the tube and on the corresponding slide was initiated at the same time. (D) Representative MFE traces at selected time points during the experiment shown in (C). Measurements were performed in different, previously unilluminated slide regions to avoid MFE attenuation caused by prior illumination. (E) Time course of the MFE amplitude during the experiment shown in (C). (F) Total autofluorescence intensity of *E. coli* BL21 cell populations (OD₆₀₀ = 8.1) in PBS with or without 1% glucose at 60-65 and 120-125 min of incubation. Two independent cultures (Day 1 and Day 2) were each measured in duplicate per condition and time window. Points show individual measurements, colored by culture. Bars and error bars indicate the mean and SD of all four measurements. Iex = 0.06 kW cm⁻². (G) Time course of extracellular pH in *E. coli* BL21 cell suspensions (OD₆₀₀ = 9.6) incubated in PBS with or without 1% glucose. Cells from a single culture were divided into two aliquots, one for each condition. (H) Glycerol dependence of the autofluorescence MFE. MFE amplitudes were evaluated over the 10–40 s irradiation window in PBS alone or PBS supplemented with 1% glucose or 1% glycerol. Each point represents the mean amplitude from one technical replicate (n = 3 per condition). Bars and error bars indicate the mean and SD across the three replicate means.

To examine the time-dependent relationship between glucose availability and MFE suppression, we monitored the MFE during sequential medium exchanges from LB to PBS, then to PBS supplemented with 1% glucose, and finally back to PBS (**Fig. 3C**). After each exchange, an aliquot was taken for slide preparation, while the remaining suspension continued through the sequence in the tube. At each stage, incubation at room temperature was initiated simultaneously for the tube suspension and the corresponding slide. MFE measurements were performed at selected incubation times in different, previously unilluminated regions of each slide. This design allowed us to track the MFE response to glucose addition and subsequent removal within a single experimental sequence while maintaining comparable cell densities across time points in both the tube suspensions and the corresponding imaging slides.

**Fig. 3D** shows representative MFE traces at selected time points during the glucose addition/removal experiment at an excitation intensity of 0.06 kW/cm². A clear negative MFE was observed in PBS, whereas the MFE was strongly suppressed after glucose supplementation. After glucose removal, the MFE remained low at 60 min but had partially recovered by 120 min. Time-course analysis of the MFE amplitude further showed that an MFE of approximately 10% was maintained in PBS, whereas the MFE remained suppressed for 120 min after glucose supplementation (**Fig. 3E**). Following glucose removal, the MFE remained low at 15–60 min and then showed progressive recovery at 90, 120, and 140 min. A similar temporal pattern was observed when the same sample was measured at a different excitation intensity (**Fig. S10-3**), making an excitation-specific origin less likely. A similar response was also observed at a higher glucose concentration (**Fig. S10-4,5**), suggesting that this behavior was not restricted to a single glucose concentration. Together, these observations indicate that the MFE is not determined solely by the immediate presence or absence of glucose in the medium. Instead, the delayed recovery is consistent with a cellular physiological and/or metabolic state that changes over time following glucose addition and removal.

We also examined whether glucose supplementation altered the cellular metabolic state under the MFE measurement conditions. Glucose supplementation decreased the total autofluorescence intensity compared with PBS alone (**Fig. 3F**), consistent with a change in the flavin-related autofluorescence/redox state^20,27,30^. In parallel, the extracellular pH remained nearly constant in PBS but progressively decreased in PBS supplemented with glucose (**Fig. 3G**), indicating glucose-dependent acidification under the measurement conditions. Acetate production was further detected by enzymatic assay under glucose-supplemented conditions (**Fig. S12**), supporting the occurrence of glucose-dependent metabolic activity. Because the MFE was also suppressed in M9 + glucose, where medium acidification was minimized by buffering (**Fig. S11 B**), extracellular acidification alone is unlikely to explain the glucose-dependent suppression of the MFE. To further examine whether the suppression of the MFE was specific to glucose-associated metabolism, we measured the MFE in PBS supplemented with glycerol as an alternative carbon source. The MFE was also strongly suppressed under the glycerol condition (**Fig. 3H**), despite the absence of the pronounced extracellular acidification observed with glucose (**Fig. S11 C**). These results further indicate that MFE suppression cannot be explained solely by glucose-associated acidification and suggest that it may instead be linked to a physiological and/or metabolic state shared between glucose and glycerol utilization.

Taken together, these results indicate that glucose-containing conditions strongly suppress the autofluorescence MFE in *E. coli* and that the MFE recovers only after a delay following glucose removal. The observation that glycerol also suppressed the MFE further indicates that this response is not specific to glucose-associated extracellular acidification. Instead, MFE suppression appears to be associated with a physiological and/or metabolic state induced by the utilization of metabolizable carbon sources. Together with the glucose-dependent changes in autofluorescence intensity, extracellular pH, and acetate production, these findings support an association between the magnetically sensitive autofluorescence component and a carbon-source-responsive cellular state, rather than a simple change in bulk autofluorescence intensity.

## Discussion

In this study, we demonstrate that endogenous autofluorescence in *E. coli* exhibits a clear magnetic field effect (MFE) exceeding 10% with the application of a 25 mT magnetic field under specific physiological and excitation conditions. Negative MFEs are observed in both BL21 and Mach1 strains. Wavelength-resolved analysis further indicated that the MFE is associated with a flavin-like component whose spectral profile differs from that of the total autofluorescence. The MFE was strongly suppressed by glucose supplementation and showed delayed recovery after glucose removal. Suppression was also observed following glycerol supplementation. Together, these findings indicate that endogenous flavin-related MFEs in *E. coli* become manifest in a metabolic-state-dependent manner.

The origin of the observed MFE can be considered in terms of both its magnetic-field dependence and spectral characteristics. The decrease in fluorescence upon magnetic-field application and the saturating field dependence observed in the MARY measurements are consistent with a radical-pair mechanism. Furthermore, the magnetic-field-induced difference fluorescence spectrum exhibited a flavin-like profile but, unlike the total autofluorescence spectrum and the illumination-induced difference spectrum, showed no clear signal in the short-wavelength region. This spectral selectivity is difficult to explain by a simple uniform modulation of the total autofluorescence and supports the involvement of a specific flavin-related component. The slight blue shift relative to free FAD further suggests that the magnetically sensitive flavin component experiences a different molecular environment, potentially involving association with intracellular macromolecules. Control measurements of free flavins, including at high concentrations, did not reproduce the >10% MFE observed in cells (**Fig. S9**). Thus, under the conditions examined, free flavins alone cannot account for the magnitude of the cellular response. The approximately 50% fluorescence MFE reported for the engineered AsLOV2-based flavin-binding protein at 10 mT demonstrates that protein-associated flavin photochemistry can generate large magnetic-field responses^34^. Taken together, these findings support a flavin-related radical-pair origin of the observed MFE and are consistent with a photoreaction involving triplet-born radical pairs.

A central finding of this study is that the autofluorescence MFE responds strongly to carbon-source availability. The MFE was strongly suppressed following glucose addition and showed delayed recovery after glucose removal. This temporal behavior suggests that the manifestation of the MFE depends on a cellular physiological or metabolic state that evolves in response to glucose addition and removal. The extracellular acidification and acetate production observed in the presence of glucose support the occurrence of cellular metabolic activity under these conditions^35^. In addition, the decrease in total autofluorescence intensity is consistent with changes in the redox state or fluorescence properties of intracellular flavin-related components^20,27,30^. Furthermore, MFE suppression was observed in glucose-supplemented M9 medium, where acidification was limited, and under glycerol-supplemented conditions without pronounced extracellular acidification. These observations suggest that pronounced extracellular acidification is not required for MFE suppression. Together, these findings support the interpretation that the manifestation of endogenous flavin-related MFEs is associated with carbon-source-responsive physiological and metabolic states in *E. coli*.

This carbon-source dependence suggests a close link between carbon metabolism and the autofluorescence MFE. We propose a working model in which a subset of flavins associated with metabolic proteins is responsible for magnetic-field sensitivity, and preferential reduction of this population suppresses the observed MFE (**Fig. 4**; see Supplementary Discussion for details). Glucose and glycerol enter central metabolism through different pathways^36,37^, and multiple flavin-dependent enzymes participate in these pathways and the associated electron transport systems^38,39^ (**Fig. 4A**). During catalysis, flavin cofactors undergo reduction and reoxidation. Depending on the enzyme and conditions, these cofactors may remain bound throughout the catalytic cycle or undergo release and rebinding^40,41^ (**Fig. 4B**). Oxidized flavins absorb blue light, whereas their two-electron-reduced forms exhibit markedly weaker absorption in this wavelength range^20,27,31^. A shift towards reduced states could therefore decrease both their fluorescence contribution and their availability for photoinduced radical-pair formation. The decrease in autofluorescence observed following carbon-source supplementation (**Fig. 3F**) is consistent with this interpretation. However, because the MFE is defined as a fractional change in total autofluorescence, ΔF/F, a proportional decrease in the fluorescence contributions of all components would leave the relative MFE unchanged if their intrinsic MFEs remained constant. Thus, if the magnetically sensitive flavin population shifts toward reduced forms and its fluorescence contribution decreases proportionally more than those of other components, the relative contribution of magnetically insensitive components to total autofluorescence increases. This could explain both the decrease in autofluorescence intensity and the suppression of the relative MFE. This model is also consistent with the spectroscopic evidence that the magnetically sensitive component accounts for only part of the total autofluorescence (**Fig. 2G**). In the triplet-born RP scheme (**Fig. 4C**), magnetic-field-dependent singlet–triplet mixing can favor the formation of long-lived radical states over rapid recombination, decreasing the fluorescent oxidized-flavin population and producing a negative MFE. An early study using differential extraction reported that free flavins accounted for less than 4% of the intracellular flavin pool in *E. coli*^42^. The magnetically sensitive population could include flavins bound within metabolic proteins or interacting with protein surfaces (**Fig. 4D**). In either environment, photoexcited oxidized flavin could accept an electron from a nearby donor, such as a tryptophan or tyrosine residue, to form an RP. A previous systematic study examined 38 proteins predicted to be FMN- or FAD-dependent in *E. coli*^33^. Based on functional annotations in EcoCyc^43^ and UniProt^44^, we selected 11 enzymes or complexes involved in glucose and glycerol metabolism and related pathways as candidates. The carbon-source dependence of the autofluorescence MFE may therefore guide the search for flavin-dependent enzymes and reaction pathways capable of exhibiting magnetic-field sensitivity through radical-pair formation in cells.

**Figure 4.**
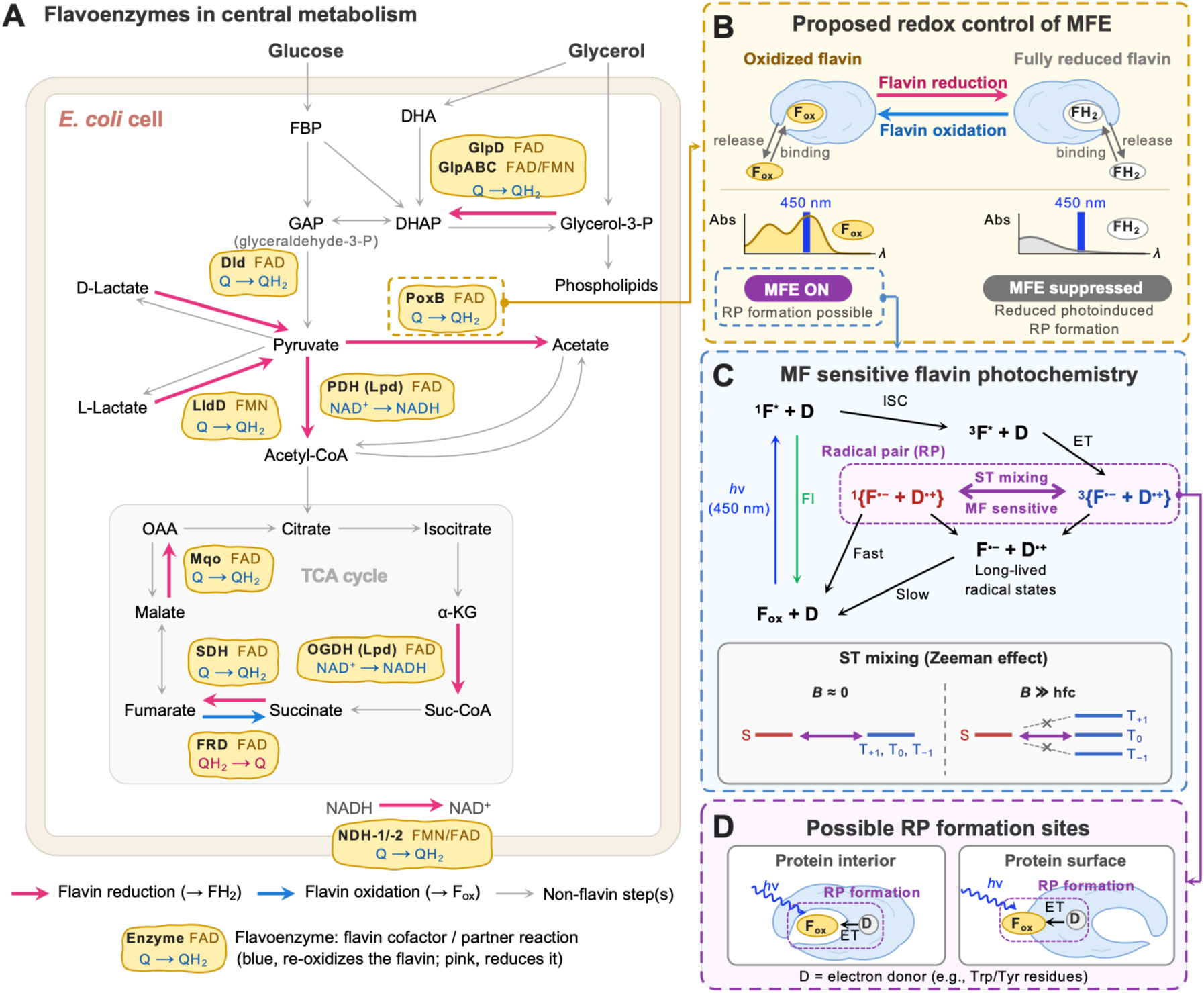
Working model for the carbon-source-dependent magnetic field effect (MFE) on flavin autofluorescence in *E. coli*. (A) Selected flavoenzymes in glucose and glycerol metabolism and related pathways, with their flavin cofactors and partner reactions (yellow boxes). Pink and blue arrows indicate steps that reduce and re-oxidize the enzyme-bound flavin, respectively; grey arrows indicate non-flavin steps. Q/QH2, oxidized/reduced quinones. (B) Proposed redox control of the MFE on flavin autofluorescence. Enzyme-bound flavin cycles between oxidized (Fₒₓ) and fully reduced (FH₂) states, with enzyme-dependent release and rebinding. Reduction decreases absorption at 450 nm and is proposed to reduce the relative contribution of MFE-active flavins to the measured autofluorescence. (C) Schematic of magnetic-field-sensitive flavin photochemistry involving triplet-born radical pairs (RPs). Excited Fₒₓ (¹F*) can fluoresce (Fl) or undergo intersystem crossing (ISC) to ³F*. Electron transfer (ET) from a donor (D) then generates a triplet RP. Magnetic-field-dependent singlet–triplet mixing modulates the competition between rapid singlet recombination and formation of long-lived radical states. In this scheme, a magnetic field can favor the latter, reducing Fₒₓ fluorescence (negative MFE). hfc, hyperfine coupling. (D) Possible RP formation sites. Photoexcited flavins within proteins or at their surfaces may form RPs by accepting electrons from nearby electron donors, such as tryptophan (Trp) or tyrosine (Tyr) residues.

The delayed recovery of the MFE magnitude after glucose removal further supports the view that the manifestation of magnetic-field sensitivity is associated with cellular states that evolve in response to nutrient availability. Links between nutritional or metabolic states and magnetic-field responses have also been reported in studies of *E. coli* under different magnetic-field conditions. Wang et al. showed that changes in NADH/NAD⁺ readouts and colony-forming unit counts following exposure to a 2.5 T pulsed magnetic field depended on prior starvation and glucose supply conditions^45^. Li et al. also reported that inhibition of colony growth by a 250 mT static magnetic field was abolished by supplementation with glycolate or glyoxylate^46^. Although the magnetic-field conditions and measured endpoints differ and a common molecular mechanism has not been established, these findings suggest that nutritional conditions and metabolic states are important considerations in understanding cellular magnetic-field responses. The autofluorescence MFE that we previously reported in HeLa cells was also observed in PBS^23^. The carbon-source dependence demonstrated here motivates investigating whether and how nutritional conditions and metabolic states during culture and measurement influence the manifestation of the MFE in HeLa cells. Future work should combine autofluorescence MFE measurements with metabolic imaging to characterize the cellular states in which magnetic-field sensitivity emerges. Furthermore, candidate flavin-dependent enzymes underlying the photochemical MFE should be investigated to determine whether radical pairs also form during catalysis in the absence of photoexcitation and whether these reactions exhibit magnetic-field sensitivity. Combining these investigations with assessments of cellular function could clarify the relationship between photochemically detected magnetic-field sensitivity, physiological redox reactions, and cellular function.

The flavin-containing molecules responsible for the observed MFE and their reaction pathways remain unidentified. Although the decrease in autofluorescence and suppression of the MFE following carbon-source supplementation are consistent with the working hypothesis of selective reduction of the magnetically sensitive flavin population, they do not directly establish this mechanism. The present data cannot distinguish a decrease in the fluorescence contribution of this population from changes in its photoreactivity, reaction branching, or intrinsic magnetic-field response. Potential contributions from changes in oxygen availability and intracellular pH associated with carbon-source utilization also require further investigation. Furthermore, steady-state MARY measurements alone cannot separate the contributions of radical-pair spin dynamics and subsequent photochemical reactions to the observed signal, and the mechanism underlying the large MFE exceeding 10% remains unresolved. Future work will combine genetic and biochemical perturbations targeting candidate molecules with assessments of flavin redox states to investigate the molecular origin of the MFE and the mechanism of its suppression by carbon sources. We also plan to combine time-resolved measurements using PP/PFP fluorescence microscopy^47^ with RYDMR and reaction kinetic modeling to examine the contributions of spin dynamics and subsequent reactions.

This study demonstrates that endogenous autofluorescence in *E. coli* exhibits an MFE with a magnitude exceeding 10% under specific physiological and excitation conditions, and that its manifestation depends strongly on carbon-source conditions. This carbon-source dependence provides a basis for investigating magnetically sensitive flavin reactions in cells in relation to their metabolic roles. A better understanding of the responsible molecules and reaction mechanisms could guide the search for naturally occurring magnetically sensitive flavoproteins beyond cryptochrome^2,4–7^, inform the design of engineered magnetically responsive proteins^34^, and support the development of label-free MFE imaging sensitive to metabolic state^48^. By combining control of metabolic state with autofluorescence MFE measurements, this approach opens a route to exploring radical-pair-derived spin effects latent within complex intracellular reaction networks and linking their manifestation to cellular states.

## Methods

### Materials

Phosphate-buffered saline (PBS) tablets (Cat. No. P4417-50TAB) and glycerol (≥99.5%; Cat. No. G9012-500ML) were purchased from Sigma-Aldrich. M9 Minimal Salts (2×; Cat. No. A1374401), LB Broth Base (Cat. No. 12780052), and D-(+)-glucose solution (45% (w/v); Cat. No. 079-05511) were purchased from Thermo Fisher Scientific, Invitrogen, and FUJIFILM Wako Pure Chemical Corporation, respectively. PBS buffer and LB medium were prepared at the appropriate concentrations using Milli-Q water and sterilized by autoclaving. M9 minimal medium was prepared at the appropriate concentration on the day of measurement using autoclaved Milli-Q water, and glucose-containing solutions were prepared by diluting the D-(+)-glucose stock solution into the corresponding medium or buffer immediately before use. PBS, PBS containing glucose or glycerol, or M9 minimal medium containing glucose was used as the imaging medium, depending on the experimental condition.

### Bacterial Strains and Cell Culture

Escherichia coli strains BL21(DE3) and Mach1 were used in this study. For fluorescence MFE and spectral measurements, a single colony grown on an LB agar plate was inoculated into 3 mL of LB medium in a sterile polypropylene culture tube (WATSON, Cat. No. 2332-004S) and cultured at 30°C for 20–24 h with continuous shaking at 140 rpm. For viability, extracellular pH, and acetate measurements, cells were cultured under the same conditions for 20 h unless otherwise stated.

### Sample Preparation for Fluorescence Measurements

The schematic protocol is shown in Fig. 1A. After shaking culture, a 0.5- to 0.8-mL aliquot of the *E. coli* culture was transferred to a 1.5-mL Eppendorf tube and centrifuged at 3,381 x g for 1 min at room temperature. After removal of the supernatant, the pellet was washed twice by resuspension in 1 mL of the imaging medium followed by centrifugation under the same conditions. After the final wash, the pellet was resuspended in the imaging medium. Before slide preparation, an aliquot of the resuspended cell suspension was diluted for OD600 measurement, and the cell density was adjusted to the desired OD600. For slide preparation, a 0.12-mm-thick SecureSeal imaging spacer (Sigma-Aldrich, Cat. No. GBL654004) was attached to a glass coverslip (No. 1, 24 × 60 mm, 0.13–0.17 mm thick; Matsunami). A 5-μL aliquot of the cell suspension was spotted at the center of the spacer and sandwiched with another glass coverslip of the same type to prepare the imaging sample.

### Fluorescence microscopy setup

The fluorescence microscope used in this study was a custom-built wide-field system based on the optical arrangement described previously^23^, with modifications to the laser and fiber-coupling optics. A 449-nm laser beam from a CUBE laser (Coherent) was passed through a 105-μm-diameter multimode fiber (M43L05, Thorlabs), collimated with an aspheric lens (CFC11P-A, f = 11 mm; Thorlabs), and filtered using a band-pass filter (AT450/50, Chroma). The excitation beam was reflected by a dichroic mirror (T470lpxr, Chroma) and focused onto the sample through a 100× oil-immersion objective lens with a numerical aperture of 1.49 (UAPON100XOTIRF, Olympus), giving a beam diameter of 17.2 μm at the sample plane. Fluorescence from the sample was collected through the same objective lens, transmitted through the dichroic mirror, directed by a reflection mirror (PFR10-P01, Thorlabs), imaged with a tube lens (AC254-200-A, f = 200 mm; Thorlabs), passed through a 500-nm long-pass emission filter (ET500lp, Chroma), and recorded using an sCMOS camera (ORCA-Flash4.0 V3, Hamamatsu) (**Fig. S1**).

### Magnetic-field generation and modulation

Magnetic fields were generated using a projected-vector-field electromagnet (GMW 5204; GMW Associates), as described previously^23^. The electromagnet was operated under the same cooling conditions as those used in the previous study, in which no detectable thermal and mechanical artifacts during fluorescence measurements were observed^23^. As in the previous study, the electromagnet was mounted above the sample and positioned within 5 mm of the imaging sample. The magnetic-field magnitude was calibrated before measurements using a high-resolution three-dimensional Hall-effect sensor (F3A, SENIS) placed in contact with the objective lens under the same magnet–objective distance used during imaging. For fluorescence MFE and MARY measurements, vertical magnetic fields relative to the sample plane were generated by applying equal currents to all three coils.

### Autofluorescence MFE Measurements

For autofluorescence MFE measurements, the focal plane was first adjusted under bright-field illumination to avoid photobleaching of autofluorescence before blue-light excitation. *E. coli* cells were then imaged under continuous-wave (CW) irradiation with 449-nm laser light. During standard MFE measurements, the applied vertical magnetic field was square-wave modulated between 0 and 25 mT at the frequency specified for each experiment. For MARY measurements, the vertical magnetic field was triangle-wave modulated between +29.25 and −29.25 mT at 0.05 Hz. The laser power was monitored using a laser power meter (LP-1, Sanwa), and the irradiation intensity was estimated from the measured laser power and the irradiation spot area calculated from the spot diameter. All fluorescence images were acquired with a camera exposure time of 100 ms.

### Fluorescence spectrum measurements

Fluorescence spectra were recorded using a CCD-based mini-spectrometer (C10083CA, Hamamatsu). The focal plane was first adjusted under bright-field illumination to avoid photobleaching of autofluorescence before blue-light excitation. The detection path was then switched from the sCMOS camera to the spectrometer by rotating the reflection mirror in the detection optics (**Fig. S1**). Spectral measurements were performed under continuous 449-nm excitation, with an excitation intensity of 0.5 kW/cm² at the sample. Background spectra were acquired under the same optical configuration and subtracted from the sample spectra during spectral analysis.

### Image/data analysis and MFE extraction

Image analysis was performed using ImageJ. For each fluorescence image frame, the average fluorescence intensity was calculated within a region of interest (ROI) covering the illuminated bacterial population. The resulting fluorescence time traces were analyzed using custom-written Python scripts and Igor Pro. To remove the magnetic-field-independent photobleaching component associated with blue-light irradiation and reveal the modulated magnetic-field-dependent component, the fluorescence decay was fitted with an offset double-exponential function using least-squares curve fitting over the selected fitting interval. Normalized residual traces were then calculated from the deviation of the observed fluorescence intensity from the fitted decay curve relative to the fitted intensity. Details of the fitting procedure, MFE extraction, and MARY analysis are provided in the Supplementary Information.

### Viability Assay (Colony Forming Units)

For viability assays, *E. coli* cells were collected by centrifugation at 3,320 × g for 1 min using a Microfuge 20R (Beckman Coulter). The cell pellets were washed twice with PBS and resuspended in PBS. The cell suspensions were maintained at room temperature. At 0, 30, 60, and 120 min after resuspension, aliquots were collected, serially diluted in PBS, and spread onto LB agar plates. After overnight incubation at 37°C, colony-forming units (CFUs) were counted, and viable cell numbers per milliliter were calculated.

### Extracellular pH Measurement Assay

For extracellular pH measurements, *E. coli* cells were harvested by centrifugation at 1,740 × g for 10 min using a swinging-bucket rotor. The cell pellets were washed twice with PBS and resuspended in either PBS alone or PBS containing 1% (w/v) glucose. After resuspension, the cell density was adjusted to the desired OD600. The suspensions were incubated at room temperature without agitation. The pH of the glucose-supplemented suspension was monitored at 3-min intervals for the first 30 min, every 5 min from 30 to 60 min, and every 15 min from 60 to 120 min. The control suspension in PBS alone was measured at 0 and 120 min. All pH measurements were performed using a calibrated HI991002 pH meter (Hanna Instruments, Woonsocket, RI, USA).

### Reproducibility and statistical analysis

The large autofluorescence MFE, the spectral characteristics of both the autofluorescence and the wavelength-resolved MFE, and MFE suppression following carbon-source supplementation were each confirmed in at least three independent experiments using separately cultured samples on different days. Definitions of technical and biological replicates for autofluorescence MFE measurements, together with details of the statistical analysis, are provided in the Supporting Information.

## Author contributions

N.I. and J.R.W. conceived and designed the study. N.I. developed and optimized the *E. coli* sample-preparation protocol, performed the fluorescence experiments and data analysis, interpreted the results, and wrote the manuscript. M.O. prepared the *E. coli* cultures, performed viable cell counting, pH and acetate measurements, interpreted the results, and wrote the relevant Methods sections. J.R.W. secured funding and supervised the study. All authors reviewed, edited and approved the manuscript.

## Acknowledgements

We thank Yan Tang for participating in initial preliminary experiments. This work was supported by JSPS KAKENHI Grant Number JP23H01919 and the U.S. National Science Foundation (NSF) Trailblazer Engineering Impact Award No. 2421670. N.I. and M.O. were supported in part by the U.S. NSF Trailblazer Engineering Impact Award No. 2421670.

