## Supporting Information for "Metabolic-State-Dependent Magnetic Field Effects on Endogenous Flavin Autofluorescence in Escherichia coli"

**Contents**

1. Microscopy setup
2. Estimation of the number of observed *E. coli* cells
3. MFE analysis
4. MARY analysis
5. MFE amplitude quantification and statistical analysis
6. Effects of experimental conditions on the autofluorescence MFE
   6.1 Excitation-intensity dependence
   6.2 Cell-density dependence
7. Assessment of background in fluorescence spectroscopy
8. Assessment of flavin leakage
9. Free FAD MFE measurement
10. Glucose addition and removal experiments
11. External pH measurement
12. Acetate measurement
13. Experimental parameters
14. Supplementary Discussion: Candidate flavin-related enzymes and metabolic pathways in *E. coli*
15. References

1. Microscope setup

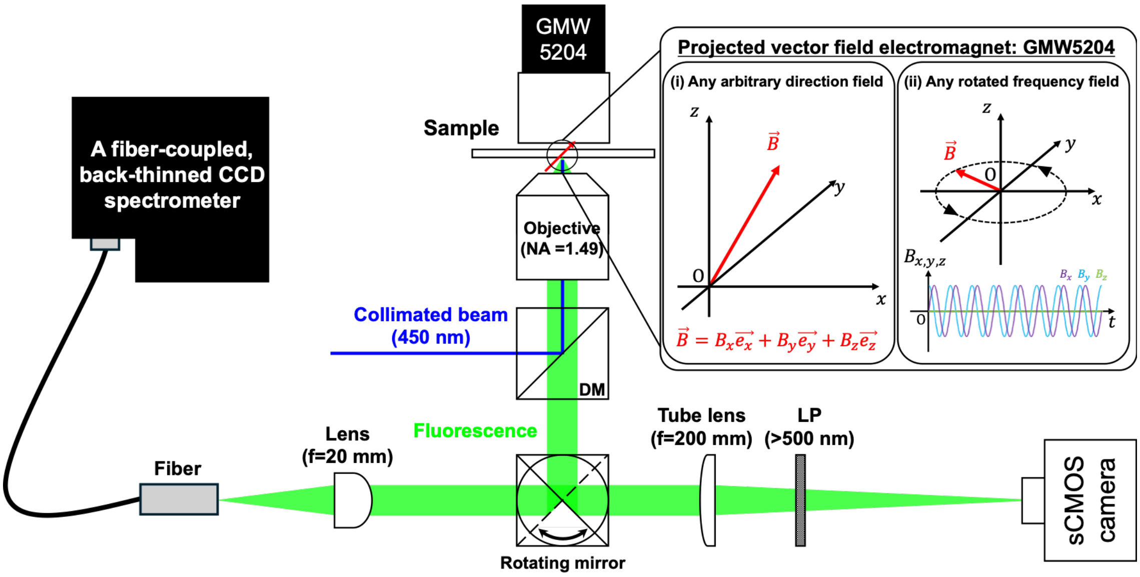


**Figure S1. Schematic of the fluorescence microspectroscopy setup used to measure magnetic-field effects on *E. coli* autofluorescence.** Fluorescence detection was switched between the sCMOS camera and the fiber-coupled, back-thinned CCD spectrometer using a rotatable mirror. DM = dichroic mirror, LP = longpass filter.

2. Estimation of the number of observed *E. coli* cells

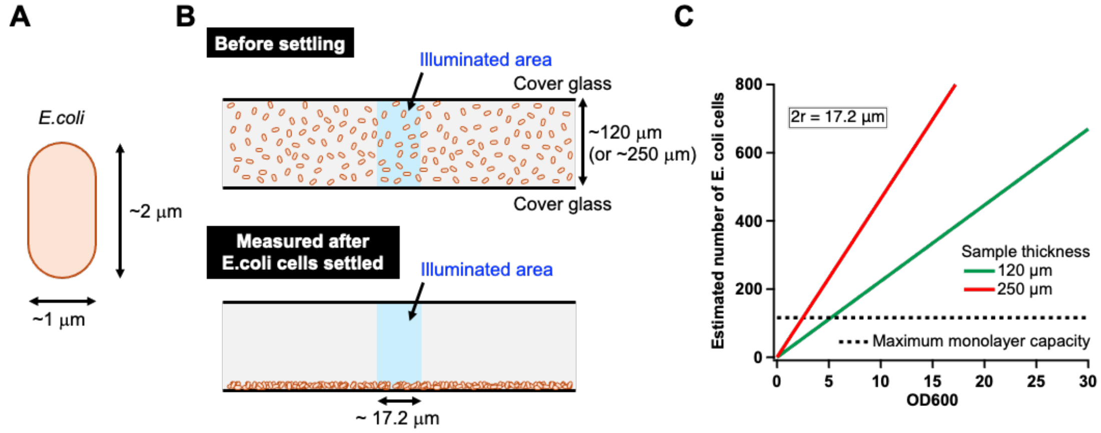


**Figure S2. Estimation of the number of *E. coli* cells within the illuminated area.** (A) Assumed representative size of an *E. coli* cell. (B) Schematic illustration of *E. coli* cell settling used to estimate the number of cells in the illuminated area. (C) Estimated number of *E. coli* cells and monolayer packing capacity in the illuminated area (diameter: 17.2 μm).

The number of *E. coli* cells contributing to the fluorescence measurement was estimated as shown in **Fig. S2**. Here, we consider a sample thickness of 120 μm. Reported *E. coli* cell dimensions are 1.6-3.1 μm in length and 0.7-1.1 μm in width[1]. For this estimate, a representative cell size of 2 μm × 1 μm was assumed, corresponding to a projected area of 2 μm² (**Fig. S2A**). Except for glucose-switch experiments, cells were allowed to settle on the glass bottom for at least 30 min before measurement. We therefore assumed that cells within the settling-column volume above the illuminated area had settled onto the glass bottom before measurement (**Fig. S2B**).

The settling-column volume was calculated as

$$V_{\mathrm{settle}} = \pi r^2 h = \pi \times (8.6\,\mu\mathrm{m})^2 \times 120\,\mu\mathrm{m} = 2.79 \times 10^{-8}\ \mathrm{mL}$$

where r is the radius of the illuminated area and h is the chamber height. Assuming that an $OD_{600}$ of 1.0 corresponds to 8.0 × $10^8$ cells $mL^{-1}$, the number of cells within this volume at $OD_{600}$ = 10 was estimated as

$$N_{\mathrm{cell}} = 10 \times 8.0 \times 10^8\ \mathrm{cells\ mL^{-1}} \times 2.79 \times 10^{-8}\ \mathrm{mL} = 223.2\ \mathrm{cells}$$

The monolayer packing capacity of the illuminated area was then estimated from the illuminated area and the projected area of one cell (**Fig. S2C**). For an illumination diameter of 17.2 μm, $A_{\mathrm{illum}} = \pi \times (8.6\,\mu\mathrm{m})^2 = 232.4\,\mu\mathrm{m}^2$. Thus, the maximum number of cells within the illuminated area under complete monolayer packing was

$$N_{\text{mono}} = \frac{232.4\ \mu\text{m}^2}{2.0\ \mu\text{m}^2} = 116.2\ \text{cells}$$

Thus, under the all-settling assumption, $OD_{600}$ = 10 corresponds to approximately two monolayer equivalents of *E. coli* cells in the illuminated area.

3. MFE analysis

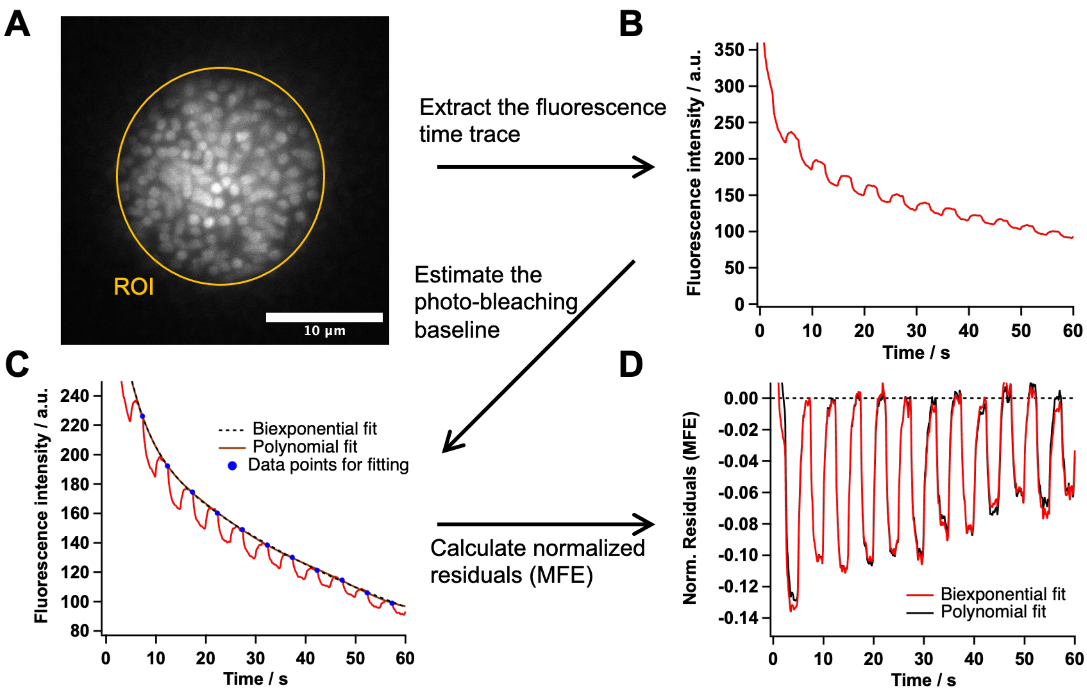


**Figure S3. Workflow for autofluorescence MFE analysis.** (A) Representative autofluorescence image with the ROI. (B) ROI-averaged fluorescence time trace. (C) Fluorescence trace with the fitted baseline curves. (D) Normalized residual traces obtained using the two baseline functions.

The workflow for autofluorescence MFE analysis is shown in **Fig. S3**. The analysis consisted of fluorescence time-trace extraction, baseline estimation, and MFE extraction.

Fluorescence time-trace extraction

For each autofluorescence image sequence, a fixed region of interest (ROI) encompassing the illuminated *E. coli* cell population was defined. The mean fluorescence intensity within the ROI was calculated for each frame. The light-induced background, measured using a PBS blank at the same excitation intensity, was subtracted to obtain the fluorescence time trace, I(t) (**Fig. S3 A,B**).

Baseline estimation and MFE extraction

The fluorescence time trace contains fluorescence decay due to photobleaching under continuous blue-light excitation, together with a decrease induced by magnetic-field application. To extract the magnetic-field-induced fluorescence change, the photobleaching decay was estimated as a baseline for subsequent correction. The first magnetic-field ON/OFF modulation cycle after excitation onset was excluded because it exhibited rapid fluorescence decay. To estimate the fluorescence intensity in the absence of a magnetic-field-induced change, the data point immediately preceding each subsequent OFF-to-ON transition was used for baseline fitting. The baseline, $I_{fit}(t)$, was estimated by least-squares fitting of the selected points using either of the following functions: a double-exponential function or a polynomial function (**Fig. S3 C**).

A double exponential function:

$$I_{fit}(t) = A_1 e^{-k_1 t} + A_2 e^{-k_2 t} + C \tag{S3-1}$$

A polynomial function:

$$I_{fit}(t) = \sum_{k=0}^{n} a_k \, t^k \tag{S3-2}$$

An eighth-order polynomial (n = 8) was used. The MFE at each time point was calculated as the normalized deviation of the observed fluorescence intensity from the fitted baseline, as follows:

$$MFE(t) = \frac{I(t) - I_{fit}(t)}{I_{fit}(t)} \tag{S3-2}$$

Here, $I_{fit}(t)$ represents the estimated fluorescence intensity in the absence of a magnetic-field-induced change. Negative MFE values indicate a decrease in fluorescence intensity upon magnetic-field application (**Fig. S3 D**). The analysis was performed using custom-written Python code.

4. MARY analysis

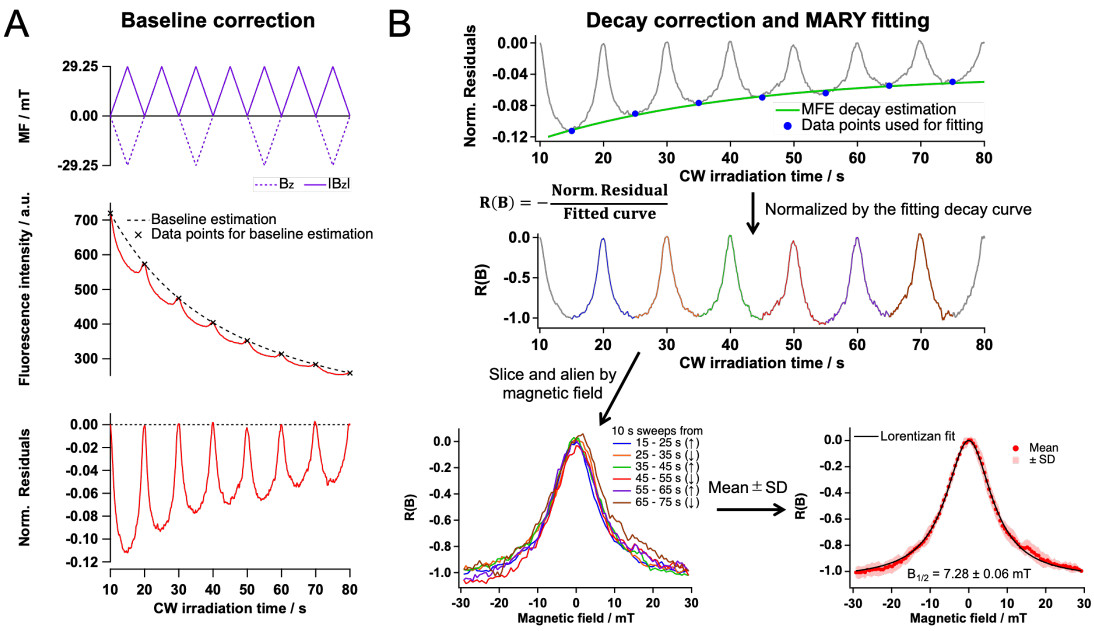


**Figure S4. Workflow for autofluorescence MARY analysis.** (A) Baseline correction. Top: applied vertical magnetic field $B_z$ (dotted) and $|B_z|$ (solid). Middle: autofluorescence intensity (red) and the baseline (black dashed line) fitted to zero-field measurements (black crosses). Bottom: normalized residuals, r(t). (B) Decay correction and MARY fitting. Normalized residuals at ($|B_z|$=29.25 mT) (blue points) were fitted to estimate the decay curve, d(t) (green). The signed normalized response was calculated as $R(B_z(t))$ = -r(t)/d(t). Six consecutive 10-s sweeps from 15 to 75 s were extracted and aligned on a common magnetic-field axis. Colors identify individual sweeps; arrows indicate sweep direction. Red points and pale red shading show the mean and ±SD across the six sweeps, respectively. The black line shows a Lorentzian fit to the mean.

The workflow for autofluorescence MARY analysis is shown in **Fig. S4**. The magnetic field was applied using triangular-wave modulation at 0.05 Hz with a peak amplitude of 29.25 mT. The fluorescence baseline was estimated from data points at zero-field crossings using the fitting procedure described in the preceding section (**Fig.S4 A**).

The saturating magnetic-field dependence associated with the radical pair mechanism (RPM) can be approximated by a Lorentzian function. For the negative MFE observed here, we used

$$MFE(B) = -MFE_{sat}\frac{B^2}{B^2 + B_{1/2}^2} \qquad \text{(S4-1)}$$

where $MFE_{sat}>0$ is the magnitude of the saturation MFE and $B_{1/2}$ is the magnetic-field magnitude at which the response reaches half its saturation magnitude.

The observed amplitude decay during illumination may reflect irreversible photochemical deactivation of magnetically responsive molecules without sufficient replenishment, reducing their relative contribution to total fluorescence. Assuming that the field-dependent line shape remains unchanged, we introduced an amplitude-decay factor, D(t):

$$MFE(B,t) = -MFE_{sat}\frac{B^2}{B^2+B_{1/2}^2}D(t) \qquad \text{(S4-2)}$$
$$D(t_0) = 1$$

Here, $t_0$ is a reference time, and $MFE_{sat}$ denotes the saturation magnitude at that time. Let $B_{pk} = max_t|B(t)|$ denote the maximum absolute field. Because Eq. S4-2 is symmetric in B, the responses at $+B_{pk}$ and $-B_{pk}$ share a common decay envelope, $MFE_{pk}(t)$. This envelope is estimated by fitting the residuals at the sweep extrema to a single-exponential function with a constant offset:

$$d(t) = a\mathbf{e}^{-k(t-t_0)} + b \qquad \text{(S4-3)}$$

The fitted curve d(t) estimates the signed response $MFE_{pk}(t)$, while $d(t)/d(t_0)$ estimates the relative decay factor D(t).

Normalization by the extremal-field response eliminates the time-dependent factor:

$$R(B) = -\frac{MFE(B,t)}{MFE_{pk}(t)} = -\frac{B_{pk}^2+B_{1/2}^2}{B_{pk}^2}\frac{B^2}{B^2+B_{1/2}^2} \propto -\frac{B^2}{B^2+B_{1/2}^2} \qquad \text{(S4-4)}$$

The minus sign preserves the negative MFE convention, giving (R(B)=-1) at ($|B|= B_{pk}$) under the model. The experimental normalized trace is calculated as R(B(t)) = -r(t)/d(t). The normalized trace is divided into individual sweeps, which are aligned on a common magnetic-field axis. For N sweeps, the mean response and sample standard deviation are

$$\bar{R}(B) = \frac{1}{N}\sum_{j=1}^{N} R_j(B)$$
$$s_R(B) = \sqrt{\frac{1}{N-1}\sum_{j=1}^{N}\left[R_j(B) - \bar{R}(B)\right]^2} \qquad \text{(S4-5)}$$

The fractional MFE scale at the reference time ($t_0$) is recovered by restoring the corresponding extremal-field amplitude:

$$MFE_{corr}(B) = -MFE_{pk}(t_0)R(B) = -MFE_{sat}\frac{B^2}{B^2+B_{1/2}^2} \qquad \text{(S4-6)}$$

When $t_0$ is chosen at a sweep extremum, the measured residual magnitude $c_0 = |r(t_0)|$ provides an estimate of $MFE_{pk}(t_0)$. The corrected mean and SD are therefore calculated as

$$\overline{MFE}_{corr}(B) = c_0\bar{R}(B), \qquad s_{crct}(B) = c_0 s_R(B) \tag{S4-7}$$

For the present data, the magnetic field was applied using triangular-wave modulation at 0.05 Hz with $B_{pk}$ = 29.25 mT. The seven residual values at t = 15,25,35,45,55,65, and 75 s were fitted to Eq. S4-3 by unweighted nonlinear least squares, with (k>0) and a and b unconstrained. Six consecutive 10 s sweeps spanning 15–75 s were extracted, aligned on a common magnetic-field axis, and averaged N = 6. The mean and SD of R(B) are shown in **Fig. S4B**. The reference time was set to $t_0$ =15 s, and the measured residual magnitude $c_0$=0.112056664 was used to restore the fractional MFE scale. The resulting decay-corrected MFE is shown in Fig. 1F. The displayed SD represents variation among the six sweeps. The mean value is fitted to the Lorentzian form Eq. S4-1.

5. MFE amplitude quantification and statistical analysis

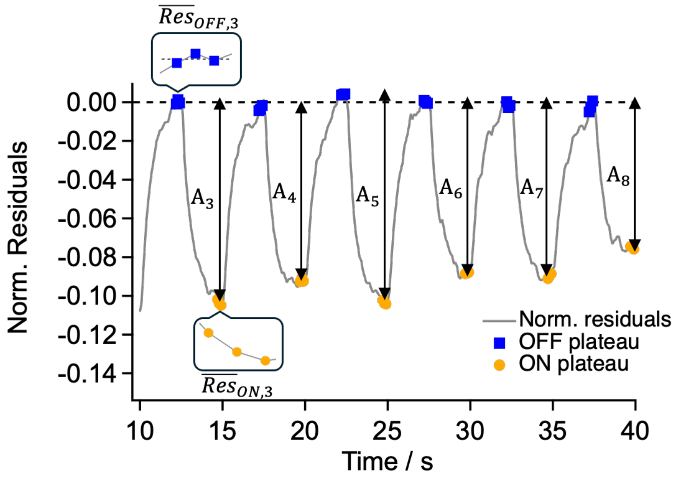


**Figure S5. Determination of MFE amplitude from normalized fluorescence residuals.** The black line shows the normalized fluorescence residuals. Blue circles and orange squares mark the data points selected near the end of each ON and OFF interval, respectively. Typically, three points were averaged per interval. The MFE amplitude for each cycle was calculated as the ON average minus the OFF average. The mean MFE amplitude and its sample standard deviation (SD) were then calculated across the analyzed cycles, typically six cycles within the 10-40 s interval.

The autofluorescence of *E. coli* exhibited a delayed response to magnetic-field ON/OFF switching. To quantify the MFE amplitude, normalized fluorescence residuals were averaged separately over several consecutive data points near the end of each ON and OFF interval (**Fig. S5**). The amplitude for each cycle was then calculated as the ON average minus the OFF average. For each measurement, the mean MFE amplitude and its sample standard deviation (SD) were calculated across the analyzed cycles. The SD was used to describe cycle-to-cycle variation, including the decrease in MFE magnitude during illumination.

The MFE amplitude for the k th cycle was defined as:

$$A_k = \left| \overline{Res}_{ON,k} - \overline{Res}_{OFF,k} \right| \quad \text{(S5-1)}$$

where $\overline{Res}_{ON,k}$ and $\overline{Res}_{OFF,k}$ denote the mean normalized fluorescence residuals over the selected points in the ON and OFF intervals, respectively.

The mean MFE amplitude was calculated as:

$$MFE_{amp,mean} = \frac{1}{N_{cycle}} \sum_{k=1}^{N_{cycle}} A_k \quad \text{(S5-2)}$$

The sample SD of the cycle amplitudes was calculated as follows:

$$MFE_{amp,SD} = \sqrt{\frac{1}{N_{cycle} - 1} \sum_{k=1}^{N_{cycle}} \left(A_k - MFE_{amp,mean}\right)^2} \tag{S5-3}$$

where $N_{cycle}$ is the number of analyzed cycles.

Technical replicates were defined as measurements performed at different illumination positions within the same sample, acquired within a few minutes of one another. Biological replicates were defined as measurements of cells cultured independently in separate tubes. Each replicate was represented by the mean MFE amplitude calculated across the analyzed cycles of a single measurement. For each replicate type, the mean MFE amplitude and sample SD were calculated as follows:

$$\bar{M}_r = \frac{1}{N_r} \sum_{i=1}^{N_r} MFE_{amp,mean}^{(i)} \tag{S5-4}$$

$$s_r = \sqrt{\frac{1}{N_r - 1} \sum_{i=1}^{N_r} \left(MFE_{amp,mean}^{(i)} - \bar{M}_r\right)^2} \tag{S5-6}$$

where r denotes technical or biological replication, and $MFE_{amp,mean}^{(i)}$ is the cycle-averaged MFE amplitude obtained from the i th replicate. $N_{tech}$ is the number of measurements at different illumination positions within the same sample, whereas $N_{bio}$ is the number of independent cultures, each contributing one cycle-averaged MFE amplitude.

6. Effects of experimental conditions on the autofluorescence MFE

This section presents measurements illustrating the effects of excitation intensity and cell density on the observed autofluorescence MFE. Excitation-intensity dependence was examined using Mach1 cells with a sample thickness of 250 μm, whereas cell-density dependence was examined using BL21 cells with a sample thickness of 100 μm at an excitation intensity of 0.06 kW/cm$^2$.

In the excitation-intensity experiments, lower excitation intensities yielded smaller MFE magnitudes but slower MFE decay during continuous illumination, whereas higher intensities yielded larger MFE magnitudes accompanied by more pronounced decay. In the cell-density experiments, both the detected autofluorescence intensity and the MFE magnitude were markedly lower at the highest cell density examined ($OD_{600}$ = 32) than at the other cell densities tested. This behavior may reflect changes in local oxygen availability or other microenvironmental factors associated with dense cell packing. Density-dependent light absorption and scattering may also influence the detected fluorescence. The contributions of these factors were not separately resolved in the present experiments.

These observations therefore highlight the importance of optimizing measurement conditions for each experimental system and matching these conditions when comparing autofluorescence MFEs.

### 6.1 Excitation-intensity dependence

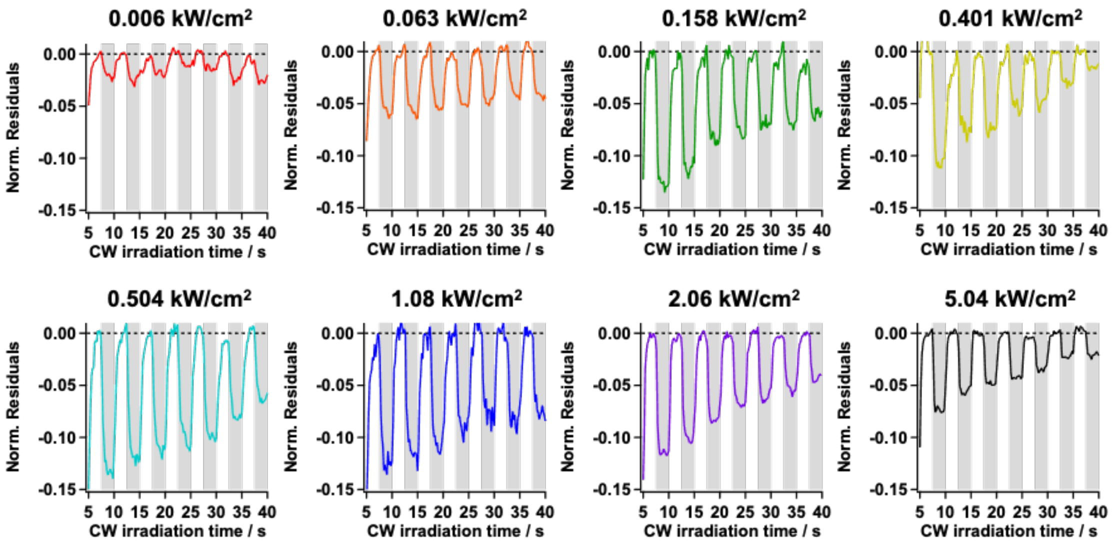


**Figure S6-1. Excitation intensity dependence of the autofluorescence MFE.** Autofluorescence MFE time traces of *E. coli* cells (Mach1) recorded under continuous-wave (CW) excitation at the intensities indicated in each panel. Measurements were performed with a sample thickness of 250 µm under ON/OFF modulation of a 25 mT magnetic field at 0.2 Hz. The traces are expressed as normalized fluorescence residuals. Gray shaded intervals indicate magnetic-field ON periods (25 mT).

6.2 Cell-density dependence of the autofluorescence MFE

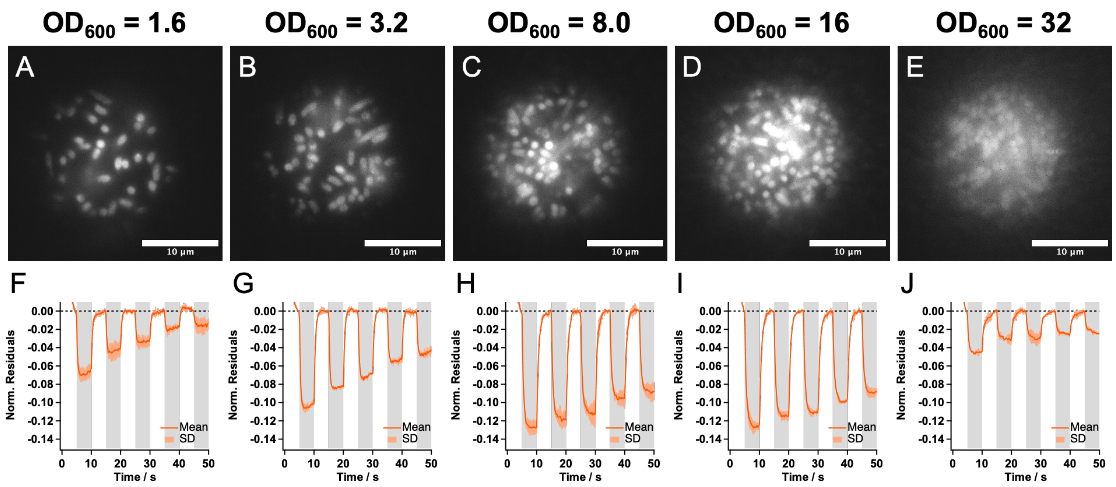


**Figure S6-2. Cell-density dependence of the autofluorescence MFE.** (A–E) Representative fluorescence images of *E. coli* cells (BL21) at $OD_{600}$ = 1.6, 3.2, 8.0, 16, and 32, respectively. All images are displayed using the same intensity scale. Scale bars, 10 µm. (F–J) Corresponding MFE time traces recorded at an excitation intensity at $I_{ex}$ = 0.06 kW/cm$^2$ after 90 min of incubation in PBS, expressed as normalized fluorescence residuals. Solid lines and shaded bands indicate the mean and ±SD, respectively, calculated at each time point across six measurements from two independent cultures, with three technical replicates per culture. At the highest cell density ($OD_{600}$ = 32), both the fluorescence intensity and the MFE magnitude were lower than those at the other cell densities examined. Gray shaded intervals indicate magnetic-field ON periods (25 mT).

7. Assessment of background in fluorescence spectroscopy

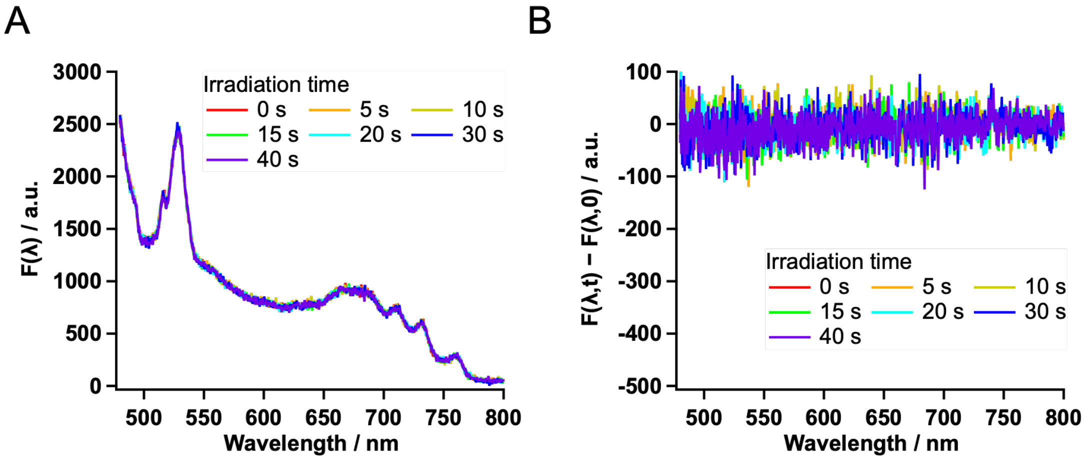


**Figure S7. Fluorescence spectra of phosphate-buffered saline (PBS).** (A) Total fluorescence spectra recorded at the indicated irradiation times under continuous 450 nm excitation at 0.5 kW/cm². (B) Corresponding illumination-induced difference spectra, F(λ,t) − F(λ,0), where t denotes irradiation time. Colors indicate irradiation times. Measurements were performed with a sample thickness of 250 µm and an integration time of 500 ms.

8. Assessment of flavin leakage

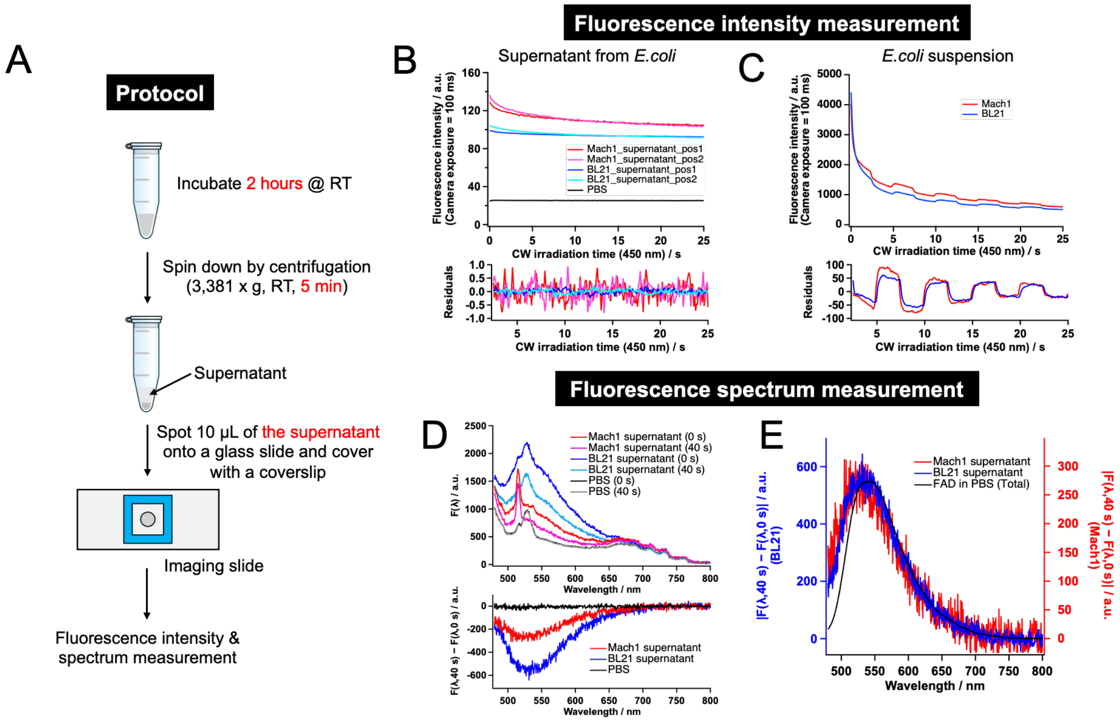


**Figure S8. Assessment of flavin leakage from *E. coli* cells.** (A) Schematic protocol for supernatant preparation and fluorescence measurements. PBS suspensions of BL21 and Mach1 cells were incubated at room temperature for 2 h and centrifuged at 3,381 × g for 5 min. A 10 µL aliquot of each supernatant was placed on a glass slide and covered with a coverslip. (B, C) Fluorescence intensity traces (upper) and corresponding residuals by biexponential fitting (lower) of the supernatants (B) and cell suspensions (C), recorded under continuous 450 nm excitation with ON/OFF modulation of a 25 mT magnetic field. PBS was measured as a background control in (B). (D) Total fluorescence spectra recorded at 0 and 40 s of irradiation (upper) and corresponding illumination-induced difference spectra, F(λ,40 s) − F(λ,0) (lower), for the supernatants and a PBS control. (E) Comparison of the spectral profiles of the sign-inverted illumination-induced difference spectra of the supernatants with the total fluorescence spectrum of free FAD in PBS. The BL21 and Mach1 supernatant signals are plotted against the left and right vertical axes, respectively.

We examined whether extracellular flavins released from *E. coli* cells could account for the MFE and fluorescence spectra observed in the cell suspensions. PBS suspensions of *E. coli* cells were prepared using the same sample preparation protocol as for all MFE measurements (Fig. 1A), up to the stage before slide preparation. The suspensions were incubated at room temperature for 2 h. Following incubation, the cells were pelleted by centrifugation at 3,381 × g for 5 min at room temperature, and the supernatants were collected. A 10 μL aliquot of each supernatant was placed on a glass slide and covered with a coverslip for fluorescence measurements (**Fig. S8A**).

Fluorescence intensity traces of supernatants from BL21 and Mach1 cell suspensions were recorded under continuous 450 nm excitation with ON/OFF modulation of a 25 mT magnetic field. Both supernatants showed fluorescence signals above the PBS background, but no clear MFE was detected in the corresponding residuals (**Fig. S8B**). Cell suspensions of both strains showed substantially stronger fluorescence signals and clear magnetic-field-dependent responses (**Fig. S8C**).

Total fluorescence spectra of the supernatants and a PBS control were recorded at 0 and 40 s of 450 nm irradiation (**Fig. S8D**, upper). To examine illumination-induced spectral changes, difference spectra, $F(\lambda, 40\ s) - F(\lambda, 0)$, were calculated (**Fig. S8D**, lower). This subtraction cancels time-invariant background contributions within the same sample, reducing sensitivity to background differences between the supernatant and an independently measured PBS control. The difference spectra revealed fluorescence loss in both supernatants, whereas the PBS control showed little change.

For comparison of spectral shape, the signs of the supernatant difference spectra were inverted, and the resulting profiles were compared with the total fluorescence spectrum of free FAD in PBS (**Fig. S8E**). Both supernatants exhibited broad spectral profiles resembling that of free FAD, consistent with the presence of extracellular flavin-related fluorescent components. This observation is consistent with previous reports of extracellular flavins in *E. coli* cultures[2,3] .

Together, the relatively weak fluorescence and absence of a clear MFE in the supernatants, compared with the stronger fluorescence and clear responses in the cell suspensions, support a predominantly cell-associated origin of the observed MFE.

9. Free FAD MFE measurement

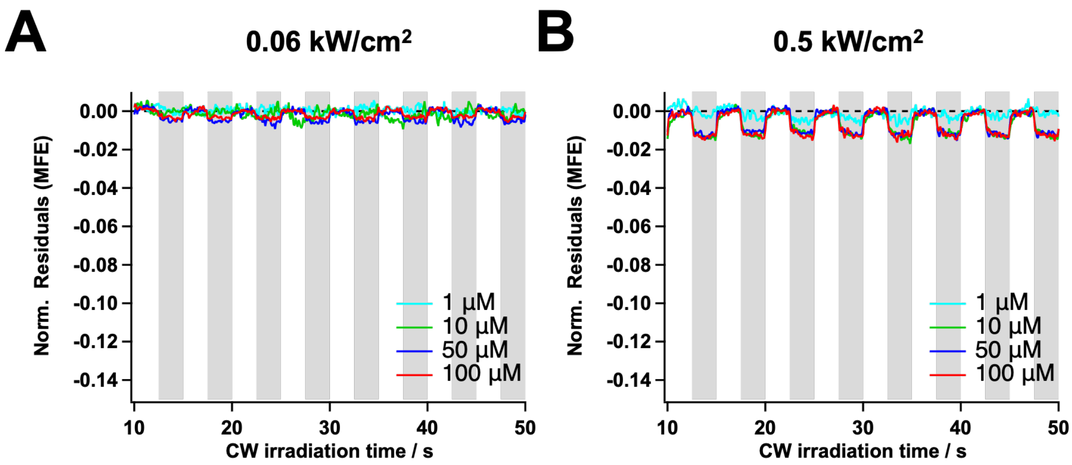


**Figure S9. Magnetic field effects on free FAD fluorescence in PBS.** Continuous-wave excitation intensities were (A) 0.06 and (B) 0.5 kW/ cm². Traces show time courses of normalized fluorescence residuals (MFE) for FAD at concentrations of 1 µM (cyan), 10 µM (green), 50 µM (blue), and 100 µM (red). Gray shading indicates magnetic-field-on periods (25 mT).

10. Glucose addition and removal experiments

This section presents the complete set of MFE time traces for the glucose addition and removal experiment shown in Fig. 3C, supplementing the representative traces in Fig. 3D. Additional measurements at a higher excitation intensity and a higher glucose concentration are also presented.

The experiments were performed with 1% and 4% glucose, each at excitation intensities of 0.06 and 0.5 kW/cm². The incubation periods and measurement times are summarized in **Fig. S10-1**. The data shown in Fig. 3D, E correspond to the 1% glucose experiment outlined in **Fig. S10A** and were acquired at the lower excitation intensity of 0.06 kW/cm², at which MFE decay during irradiation was less pronounced.

The full MFE time traces for the condition used in Fig. 3D, E are presented in **Fig. S10-2**, including measurements at the remaining incubation time points not shown in Fig. 3D. **Fig. S10-3** presents the full MFE time traces and corresponding amplitudes for the 1% glucose addition and removal experiment at 0.5 kW/cm², obtained from the same samples used for Fig. 3D, E. The full MFE time traces and corresponding amplitudes for the 4% glucose addition and removal experiments at 0.06 and 0.5 kW/cm² are presented in **Fig. S10-4** and **Fig. S10-5**, respectively. A qualitatively similar pattern of MFE suppression after glucose addition and delayed recovery after glucose removal was observed at both excitation intensities and both glucose concentrations.

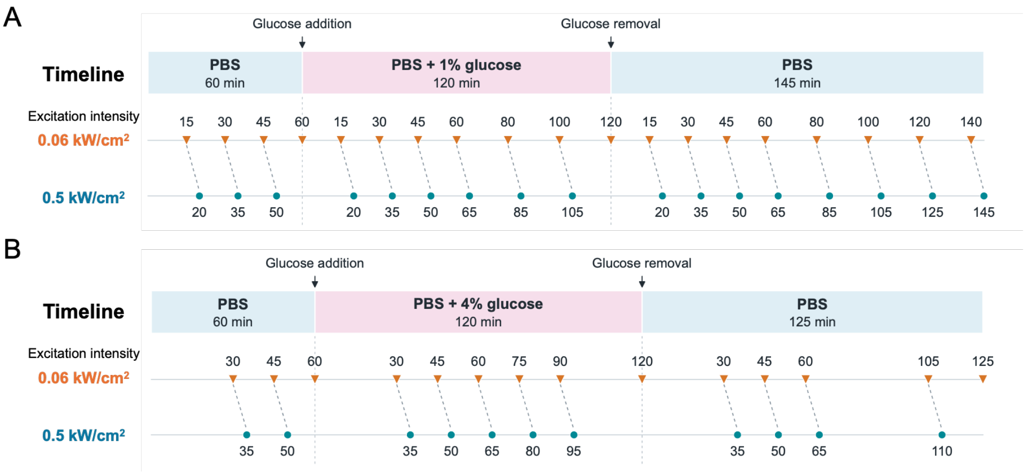


**Figure S10-1. Timelines of glucose addition and removal and MFE measurements.** (A, B) Experiments with 1% and 4% (w/v) glucose, respectively. Colored bars indicate the medium composition and duration of each incubation stage. Orange triangles and teal circles mark measurements at excitation intensities of 0.06 and 0.5 kW/cm², respectively. Numbers beside the symbols indicate incubation times in minutes from the start of each stage. Measurements were performed at different, previously unilluminated slide positions.

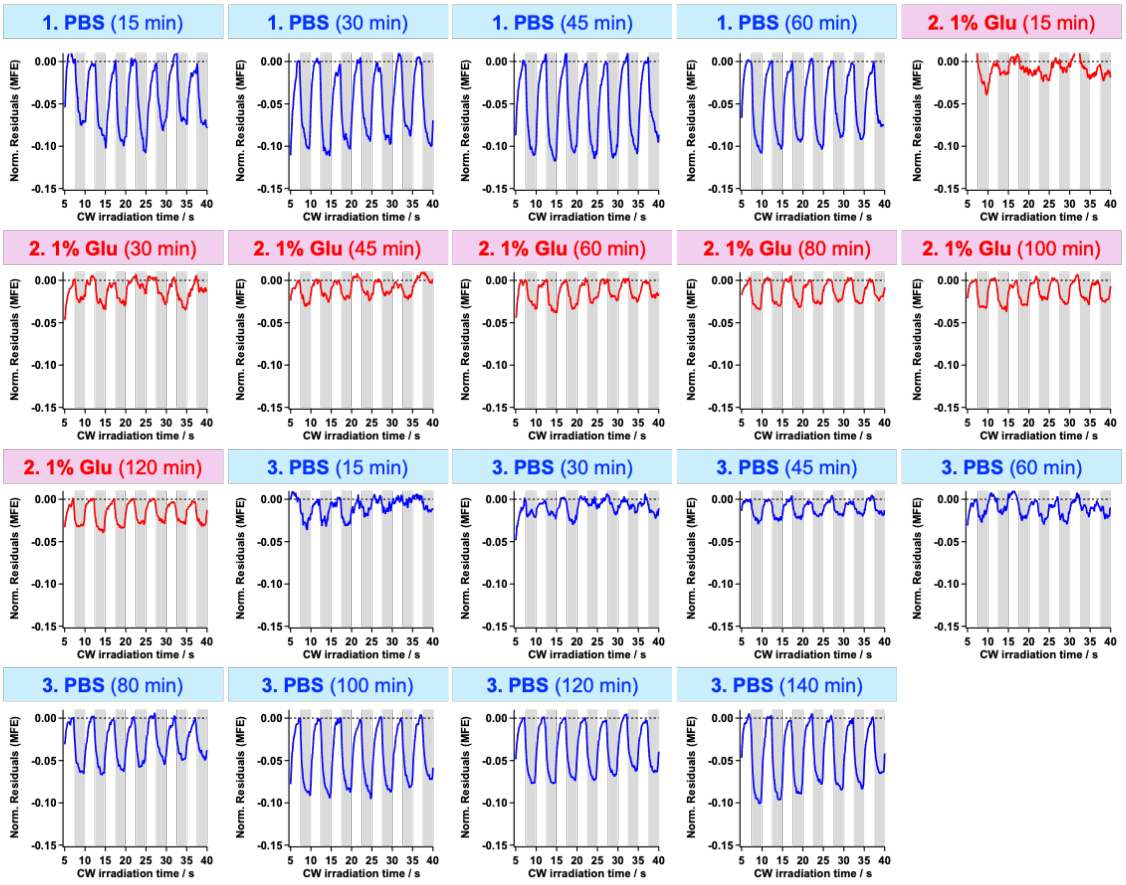


**Figure S10-2. MFE time traces during addition and removal of 1% glucose at 0.06 kW/cm².** Complete set of MFE time traces for the experiment shown in Fig. 3D, E. The numbered stages indicate PBS before glucose addition (1), PBS supplemented with 1% (w/v) glucose (2), and PBS after glucose removal (3). Blue and red traces correspond to PBS and glucose-supplemented PBS, respectively. Times in parentheses indicate incubation times from the start of each stage. Traces are expressed as normalized fluorescence residuals and plotted against irradiation time under continuous-wave excitation. Gray shaded intervals indicate magnetic-field ON periods (25 mT). Glu, glucose.

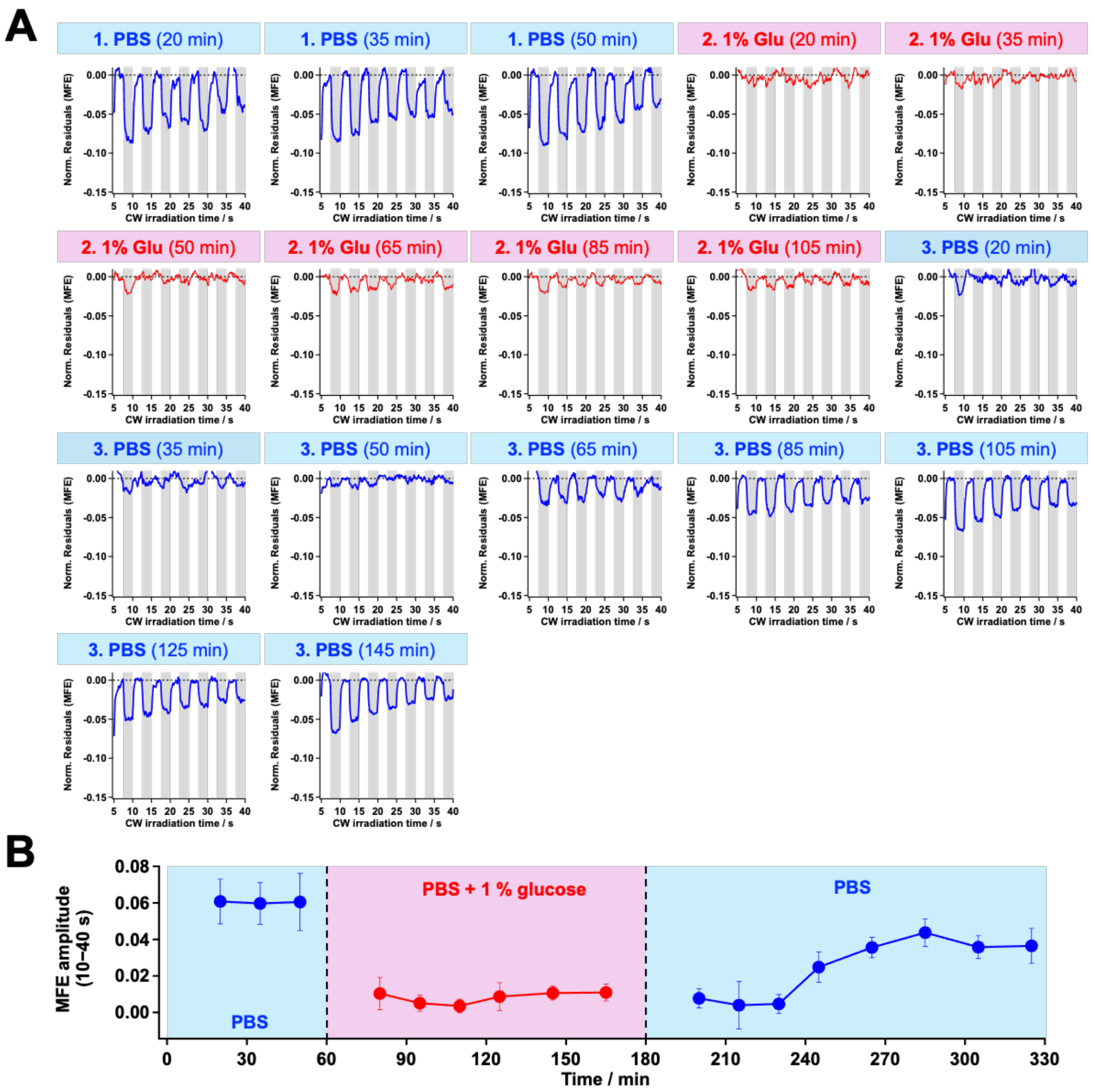


**Figure S10-3. MFE time traces and magnitudes during addition and removal of 1% glucose at 0.5 kW/cm².** (A) Complete set of MFE time traces obtained from the same samples used for Fig. 3D, E. Plotting conventions and annotations are as in **Fig. S10-2**. (B) Time course of the MFE magnitude, calculated from the cycles within the 10–40 s irradiation interval. MFE amplitudes were calculated as described in Section 5. The cycle-averaged negative MFE amplitude is displayed as a positive magnitude, with error bars indicating the sample SD of the cycle amplitudes (n = 6 cycles). The horizontal axis shows cumulative incubation time from the start of the initial PBS stage.

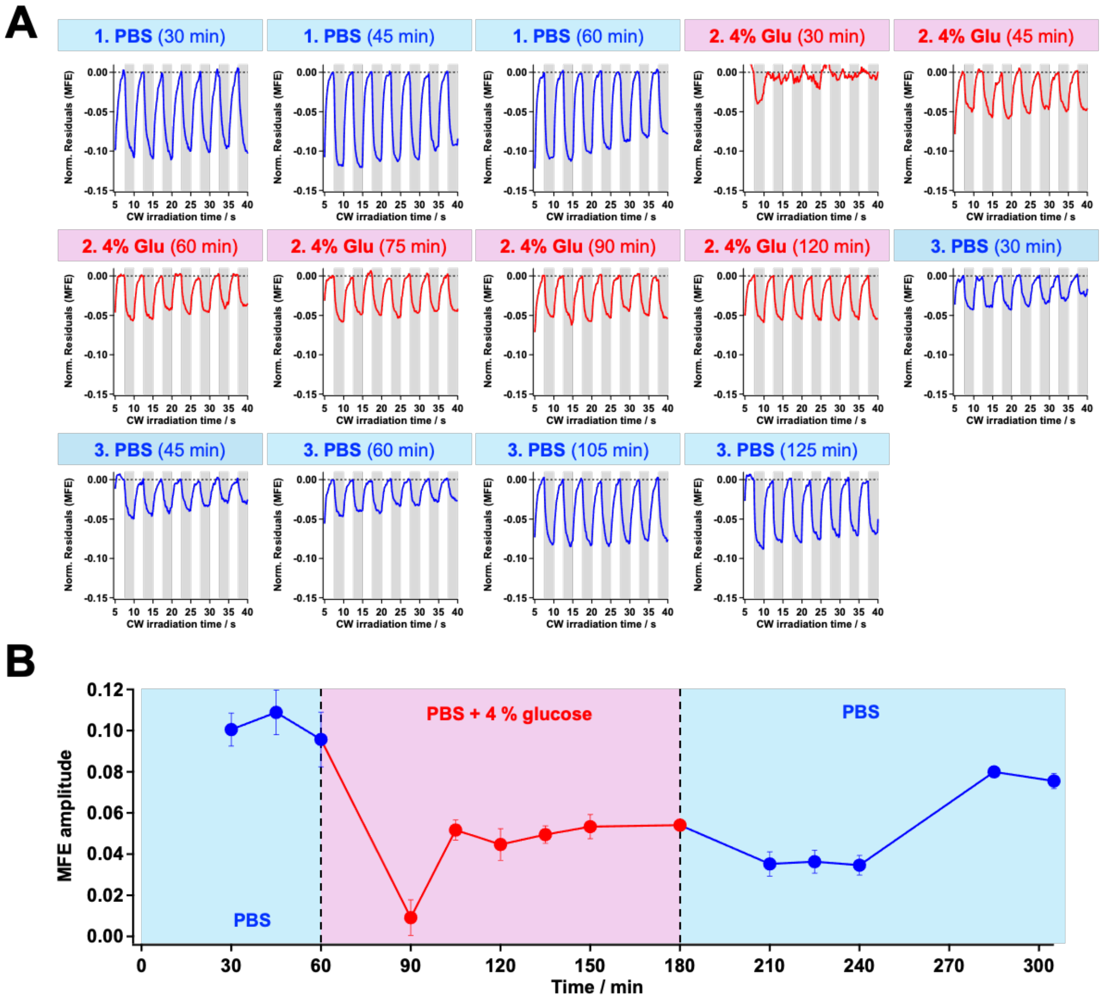


**Figure S10-4. MFE time traces and magnitudes during addition and removal of 4% glucose at 0.06 kW/cm².** (A) Complete set of MFE time traces for the 4% (w/v) glucose experiment outlined in **Fig. S10-1B**. Plotting conventions and annotations are as in **Fig. S10-2**. (B) Time course of the corresponding MFE magnitude. MFE amplitudes were calculated as described in Section 5. The cycle-averaged negative MFE amplitude is displayed as a positive magnitude, with error bars indicating the sample SD of the cycle amplitudes (n = 6 cycles). The horizontal axis shows cumulative incubation time from the start of the initial PBS stage.

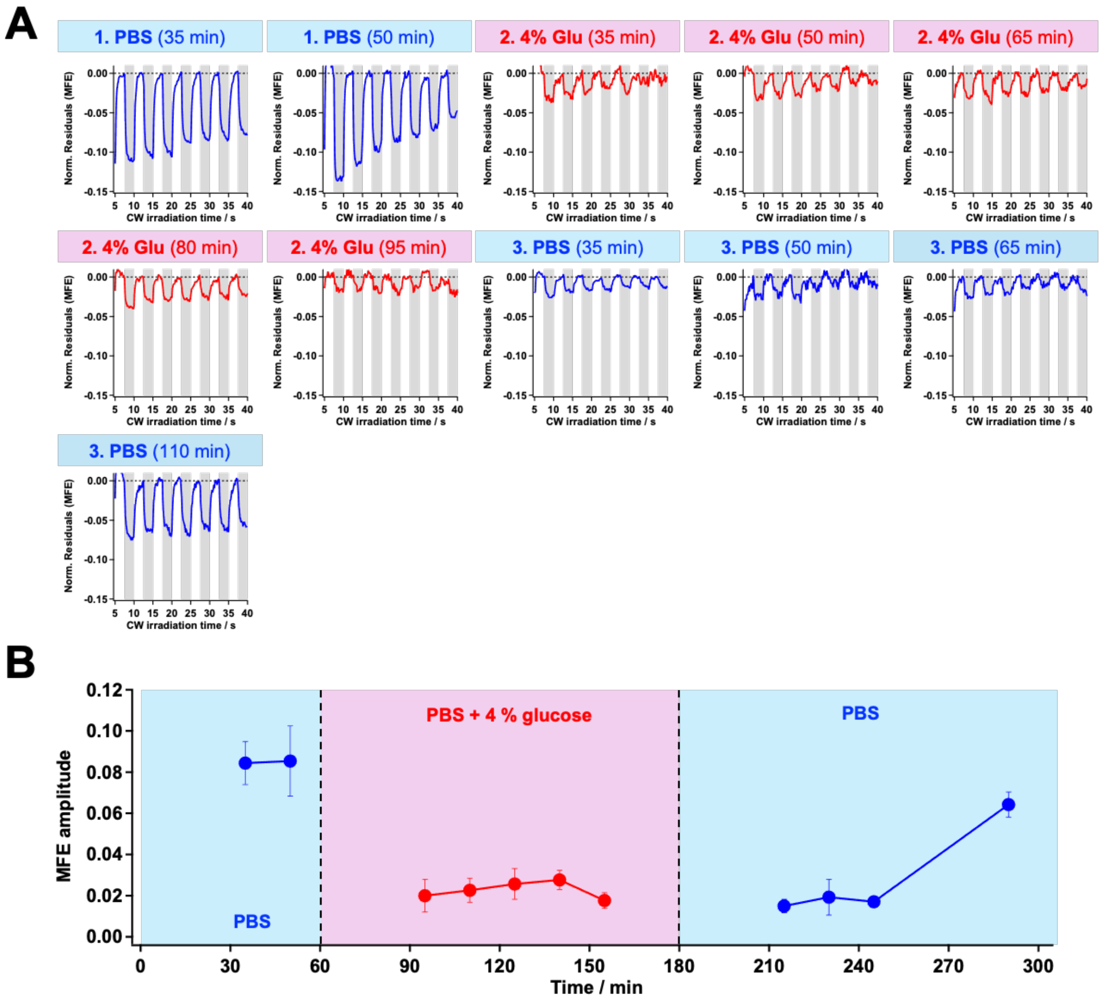


**Figure S10-5. MFE time traces and magnitudes during addition and removal of 4% glucose at 0.5 kW/cm².** (A) Complete set of MFE time traces for the 4% (w/v) glucose experiment outlined in **Fig. S10-1B**. Plotting conventions and annotations are as in **Fig. S10-2**. (B) Time course of the corresponding MFE magnitude. MFE amplitudes were calculated as described in Section 5. The cycle-averaged negative MFE amplitude is displayed as a positive magnitude, with error bars indicating the sample SD of the cycle amplitudes (n = 6 cycles). The horizontal axis shows cumulative incubation time from the start of the initial PBS stage.

11. External pH measurement

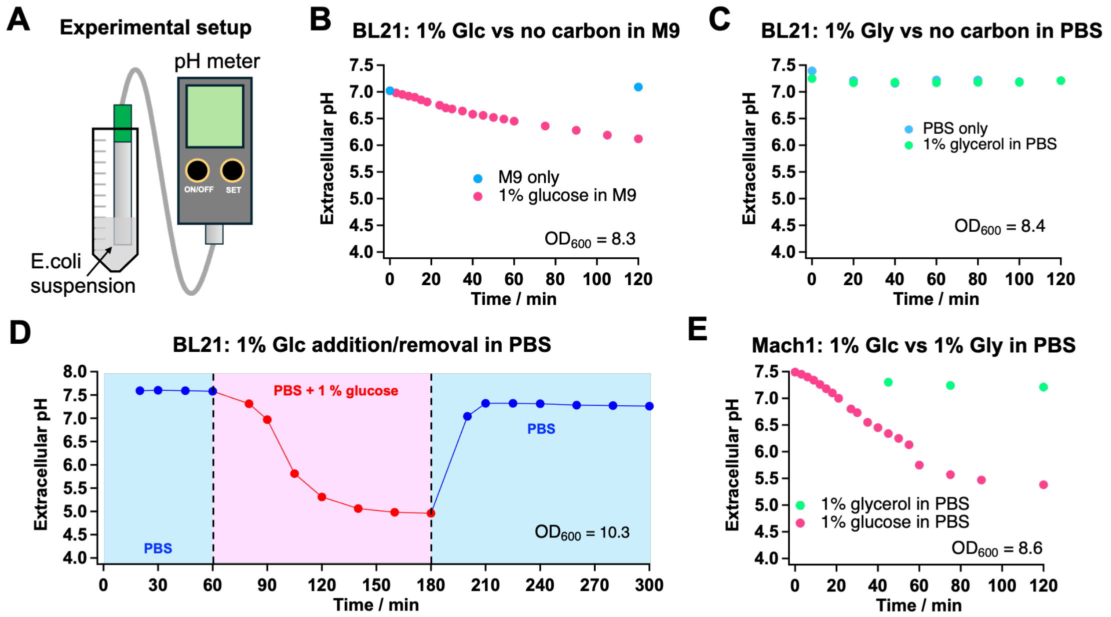


**Figure S11. Extracellular pH changes under different medium and carbon-source conditions.** (A) Experimental setup for extracellular pH measurements. (B) Extracellular pH changes in BL21 cell suspensions in M9 medium supplemented with 1% (w/v) glucose or without an added carbon source. (C) Extracellular pH changes in BL21 cell suspensions in PBS supplemented with 1% (w/v) glucose or 1% (w/v) glycerol. (D) Extracellular pH changes during sequential addition and removal of 1% (w/v) glucose in BL21 cell suspensions in PBS. (E) Extracellular pH changes in Mach1 cell suspensions under the same medium conditions as in (C). Glc, glucose; Gly, glycerol.

12. Acetate measurement

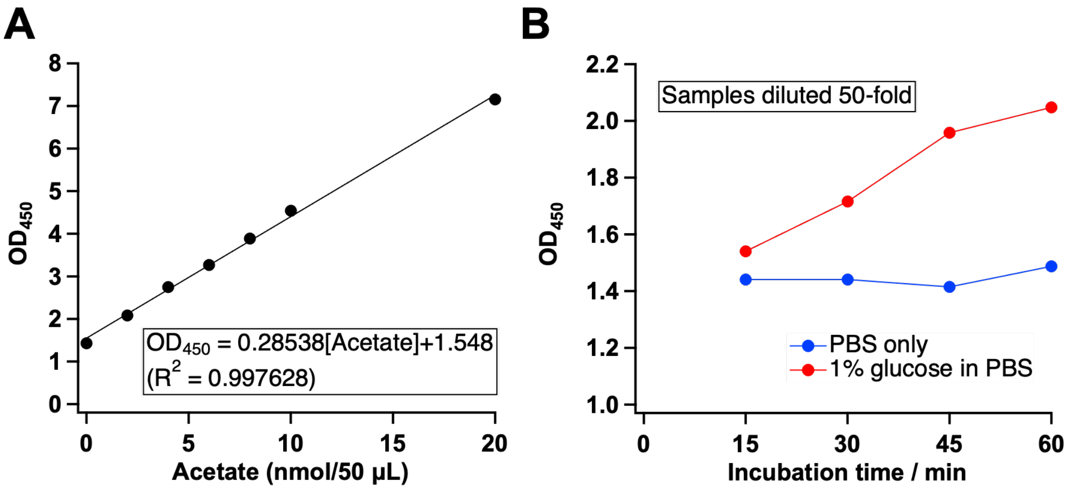


**Figure S12. Acetate assay of supernatants from *E. coli* suspensions incubated with or without glucose.** (A) Acetate standard curve obtained using 50-µL reaction mixtures. $OD_{450}$ was measured using 2-µL aliquots of the reaction mixtures. The solid line shows a linear fit. (B) Acetate assay signals ($OD_{450}$) from supernatants collected after incubation of *E. coli* cells ($OD_{600}$ = 10.3) in PBS with or without 1% (w/v) glucose for the indicated times. Supernatants were diluted 50-fold for analysis.

Acetate accumulation was qualitatively assessed using an Acetate Assay Kit (Colorimetric; ab204719, Abcam) according to the manufacturer's protocol, with minor modifications. Aliquots of the same *E. coli* cell suspensions used for extracellular pH measurements were collected after 15, 30, 45, and 60 min of incubation in PBS with or without 1% (w/v) glucose. Cells were removed by centrifugation, and the supernatants were collected and diluted 50-fold for analysis.

Reaction mixtures were prepared by combining 25 µL of the prepared sample or acetate standard solution with 25 µL of the assay reaction mix, giving a total volume of 50 µL. Following incubation at room temperature protected from light using aluminum foil, a 2-µL aliquot of each reaction mixture was loaded onto a NanoDrop OneC microvolume UV–Vis spectrophotometer (ND-ONEC-W-CP23, Thermo Fisher Scientific), and absorbance at 450 nm ($OD_{450}$) was measured. An acetate standard curve was generated using 50 µL aliquots containing 0–20 nmol acetate (**Fig. S12A**). Assay results are presented as measured $OD_{450}$ values for qualitative comparison between conditions.

The assay signals from supernatants of cells incubated in PBS without added glucose remained close to the zero-acetate standard, whereas those from glucose-supplemented suspensions increased with incubation time (**Fig. S12B**). These findings suggest that organic acid production, including acetate, may contribute to extracellular acidification.

13. Experimental parameters

**Table S1.** Sample preparation conditions for each experiment

| Figure(s) | Culture duration | $OD_{600}$ at the end of culture | $OD_{600}$ at measurement | Time from resuspension to start of measurement |
|---|---|---|---|---|
| Fig.1B | 20 h | 2.1 | 14.3 | 45 min |
| Fig.1C | 20 h | 2.1 | 14.3 | 50 min |
| Fig.1D | 21 h[a] | 3.4 (BL21)<br>3.2 (Mach1) | 15.0 (BL21)<br>15.0 (Mach1) | 30 min |
| Fig.1E | 20 h | 2.1 | 8.2 | 130 min |
| Fig.1F | 21 h[a] | 3.37 | 15.0 | 70 min |
| Fig.1G | 21 h[a] | 3.37 | 15.0 | See Fig.1G |
| Fig.2 | 20 h | 2.4 | 28.4 | 120 min |
| Fig.3A | 22 h | 2.0 | 10.0 | 60 min |
| Fig.3B | 22 h | 2.0 | 10.0 | 120 min |
| Fig.3D, E, Fig.S10-1, -2 | 24 h | 2.2 | 8.1 | See Fig.S10-1 |
| Fig.3F | 22 h | 2.0 (Day1)<br>1.6 (Day2) | 10.0 (Day1)<br>10.0 (Day2) | 60 min and 120 min |
| Fig.S6-1 | 20 h | 2.4 | 28.4 | 120 min |
| Fig.S6-2 | 22 h | 2.1 | See Fig.S6-2 | 90 min |
| Fig.S8 | 24 h | 2.4 (BL21)<br>2.2 (Mach1) | 15.0 (BL21, Fig.S8C)<br>15.0 (Mach1, Fig.S8C) | 120 min |
| Fig.S10-3, -4 | 20 h | 2.1 | 14.3 | See Fig.S10-1 |

[a] Cells were cultured in 5 mL of LB medium in a 50-mL centrifuge tube (STAR 50 ml CentrifugeTube II, cat. no. RSV-50CT2) at 140 rpm and 30°C. All other culture conditions were as described in “Bacterial Strains and Cell Culture” in the Methods section.

**Table S2.** Imaging conditions for each experiment

| Figure(s) | Sample thickness | Excitation wavelength | Excitation intensity | Illumination diameter | Camera exposure time |
|---|---|---|---|---|---|
| Fig.1, 3, S6-2, S10 | 120 μm | 450 nm | See corresponding figure legends | 17.2 μm | 100 ms |
| Fig.2, S6-1, S7, S8, S9 | 250 μm | | | | |

14. Supplementary Discussion: Candidate flavin-related enzymes and metabolic pathways in *E. coli*

This Supplementary Discussion outlines the biochemical and photochemical rationale for the working hypothesis (Fig. 4) proposed to explain the carbon-source dependence of the autofluorescence MFE. We address four points: (i) flavin redox reactions in carbon-source metabolism; (ii) flavin binding, redox state and radical-pair photochemistry; (iii) redox control of the autofluorescence MFE; and (iv) candidate flavin-dependent enzymes involved in glucose and glycerol metabolism and related pathways.

(i) Flavin redox reactions in carbon-source metabolism

*E.coli* obtains energy for cellular activities and precursors for the synthesis of cellular components by metabolizing carbon sources such as glucose and glycerol. This process involves numerous redox reactions, some of which are catalyzed by enzymes that use flavins as cofactors. In these enzymes, electrons are transferred from substrates to electron acceptors through the reduction and reoxidation of flavins. Understanding the relationship between carbon-source utilization and flavin redox reactions therefore provides a biochemical basis for examining the carbon-source dependence of the autofluorescence MFE.

Glucose and glycerol enter central carbon metabolism in *E. coli* through different routes. Glucose is metabolized to pyruvate through glycolysis, and pyruvate-derived acetyl-CoA can enter the tricarboxylic acid (TCA) cycle during respiratory metabolism. Under aerobic conditions, glycerol primarily enters glycolysis via glycerol 3-phosphate and dihydroxyacetone phosphate (DHAP). The conversion of glycerol 3-phosphate to DHAP is catalyzed by the FAD-dependent enzyme GlpD[4]. Several flavin-dependent oxidoreductases participate in these metabolic pathways and the associated respiratory chain (**Fig. 4A** and **Table S3**).

Glucose and glycerol metabolism can also differ in the balance between acetate production and reutilization. Under growth conditions in which glucose is rapidly utilized, *E. coli* can excrete acetate even under aerobic conditions through overflow metabolism[5]. In contrast, no extracellular acetate accumulation was detected in cultures of *E. coli* strain JM101 grown on glycerol under the conditions examined by Martínez-Gómez et al.[6]. To explain this observation, that study proposed that acetate produced from pyruvate via PoxB, one of the flavoenzymes shown in **Fig. 4A**, is converted to acetyl-CoA by acetyl-CoA synthetase (Acs) and reutilized through the glyoxylate shunt[6]. In our experiments with BL21 and Mach1 cells suspended in PBS, glucose supplementation decreased extracellular pH, whereas the decrease was less pronounced following glycerol supplementation (**Figs. 3G, S11C** and **S11E**).

(ii) <u>Flavin binding, redox state and radical pair photochemistry</u>

Flavin cofactors undergo reduction and reoxidation during catalysis, while their retention by the protein depends on the enzyme and conditions (**Fig. 4B**). For example, the FAD cofactors of *E. coli* succinate dehydrogenase and fumarate reductase are covalently attached to their flavoprotein subunits[7], whereas reversible FMN dissociation has been observed in NADH-reduced *E. coli* respiratory complex I[8]. Release and rebinding in **Fig. 4B** therefore apply specifically to enzymes and conditions that permit cofactor exchange.

These redox changes alter the light absorption properties of flavins. Oxidized FAD and FMN exhibit a strong absorption band near 450 nm, whereas their two-electron-reduced (hydroquinone) forms generally absorb more weakly in this spectral region[9]. Our working model proposes that preferential reduction of the magnetically sensitive flavin population decreases both its fluorescence contribution and its availability for photoinduced radical-pair (RP) formation under 450 nm excitation (**Fig. 4B**).

(iii) <u>Redox control of the autofluorescence MFE</u>

In the triplet-born RP reaction scheme shown in **Fig. 4C**, photoexcited oxidized flavin can fluoresce or undergo intersystem crossing to an excited triplet state. Electron transfer from a nearby donor then generates a triplet RP. Magnetic-field-dependent singlet–triplet mixing modulates the competition between rapid singlet recombination and formation of long-lived radical states. In this scheme, magnetic-field-induced suppression of singlet–triplet mixing can reduce the yield of rapid singlet recombination to ground-state oxidized flavin. Under illumination, this can favor the accumulation of long-lived radical states and decrease the oxidized-flavin population available for photoexcitation, producing a negative fluorescence MFE.

To examine how this affects the observed MFE, we express the detected fluorescence as the sum of a component arising from magnetically sensitive oxidized flavins and a magnetically insensitive background. The background includes magnetically insensitive oxidized flavins and other fluorescent species; the fluorescence contribution of two-electron-reduced flavins under 450 nm excitation is neglected. Let $C_{s,ox}$ and $k_s$ denote the concentration and zero-field molar fluorescence brightness of the sensitive oxidized flavins, respectively, and let $C_{ns}$ and $k_{ns}$ denote the total concentration and mean zero-field molar fluorescence brightness of the background species, respectively. The zero-field fluorescence intensities of the two components are then

$$F_s = k_s C_{s,\mathrm{ox}}, \qquad F_{\mathrm{ns}} = k_{\mathrm{ns}} C_{\mathrm{ns}} \tag{S14-1,2}$$

The background composition is assumed to remain constant, and the fluorescence brightness coefficients incorporate the excitation and detection conditions. Let $-m_s(B)$ denote the intrinsic relative MFE of the sensitive component, with $m_s(B) \geq 0$. The total fluorescence intensities at zero field and at magnetic field strength B are then

$$F(0) = F_s + F_{\mathrm{ns}}, \qquad F(B) = [1 - m_s(B)]F_s + F_{\mathrm{ns}} \tag{S14-3,4}$$

The observed relative MFE is therefore

$$\mathrm{MFE}(B) = \frac{F(B) - F(0)}{F(0)} = -m_s(B)\frac{F_s}{F_s + F_{\mathrm{ns}}} \tag{S14-5}$$

Thus, the relative MFE depends on the intrinsic MFE of the sensitive component and its fractional contribution to the total fluorescence. Below, we compare two cases assuming that $m_s(B)$ and the fluorescence brightness coefficients remain constant.

***Both components decrease by the same proportion***

If $C_{s,ox}$ and $C_{ns}$ both decrease to a fraction α ($0<\alpha<1$) of their initial values,

$$\mathrm{MFE}_{\mathrm{uniform}}(B) = -m_s(B)\frac{\alpha F_s}{\alpha F_s + \alpha F_{\mathrm{ns}}} = \mathrm{MFE}(B). \tag{S14-6}$$

The factor α cancels between the numerator and denominator, so the relative MFE remains unchanged even when the total fluorescence intensity decreases.

***Selective reduction of the magnetic field sensitive component***

If the concentration of oxidized flavins in the sensitive component decreases to a fraction α ($0\leq\alpha<1$) of its initial value while the background remains constant,

$$\mathrm{MFE}_{\mathrm{selective}}(B) = -m_s(B)\frac{\alpha F_s}{\alpha F_s + F_{\mathrm{ns}}}. \tag{S14-7}$$

For $F_s>0$, $F_{ns}>0$, and $m_s(B)>0$, the magnitude of the MFE is smaller than that before reduction.

Here, the intrinsic MFE is assumed to remain constant to isolate the effect of changes in the fluorescence contribution of the magnetically sensitive population. In the experimental system, preferential reduction of this population and changes in the intrinsic MFE arising from modulation of radical-pair spin dynamics by changes in the cellular environment may jointly contribute to the observed MFE suppression. Disentangling these contributions remains a task for future studies.

(iv) Candidate flavin-dependent enzymes involved in glucose/glycerol metabolism.

Candidates for the magnetically sensitive flavin population include flavins bound within metabolic proteins and flavins interacting with their surfaces (**Fig. 4D**). In either environment, photoexcited oxidized flavin could accept an electron from a nearby donor, such as a tryptophan or tyrosine residue, to form an RP. An early differential-extraction study reported that free flavins accounted for less than 4% of the intracellular flavin pool in *E. coli*[10]. This extraction-based estimate does not establish the flavin binding states under our measurement conditions, but provides a rationale for considering protein-associated flavins. We therefore focus on flavin-dependent proteins involved in glucose and glycerol metabolism and related pathways.

A systematic analysis of 38 proteins predicted to be FMN- or FAD-dependent in *E. coli* has been reported[11]. Based on functional annotations in EcoCyc[12] and UniProt[13], **Table S3** summarizes candidates involved in central carbon metabolism and associated pathways, including the respiratory chain, lactate utilization, and glycerol metabolism. For the 11 enzymes or complexes shown in **Fig. 4A**, the table lists their flavin cofactors, metabolic functions, and the oxidation or reduction of flavins accompanying the reactions of their respective substrates. The involvement of these candidates in the observed MFE remains to be established

**Table S3.** Flavin cofactors and metabolic functions of the candidate flavoenzymes in Fig. 4A.

| **Enzyme or complex**<br>UniProt accession(s) | **Flavin cofactor** | **Metabolic function** |
|---|---|---|
| **GlpD**<br>P13035 | FAD | Oxidizes glycerol 3-phosphate to dihydroxyacetone phosphate (DHAP) and transfers electrons to quinones during aerobic glycerol utilization.<br>Flavin reduction: FAD → $FADH_2$ |
| **GlpABC**<br>GlpA: P0A9C0<br>GlpB: P13033 | FAD/FMN[a] | Catalyzes oxidation of glycerol 3-phosphate to DHAP coupled to quinone reduction during anaerobic glycerol utilization.<br>Flavin reduction: $F_{ox}$ → $FH_2$ |
| **Lpd (PDH/OGDH)**<br>P0A9P0 | FAD | Reoxidizes reduced lipoyl groups while reducing $NAD^+$ to NADH. Serves as the shared E3 component of the pyruvate and 2-oxoglutarate dehydrogenase complexes.<br>Flavin reduction during lipoyl oxidation: FAD → $FADH_2$ |
| **PoxB**<br>P07003 | FAD | Catalyzes oxidative decarboxylation of pyruvate to acetate and $CO_2$, coupled to ubiquinone reduction.<br>Flavin reduction: FAD → $FADH_2$ |
| **SDH (SdhABCD)**<br>SdhA: P0AC41 | FAD | Couples succinate oxidation to fumarate with quinone reduction, linking the TCA cycle to respiration.<br>Flavin reduction: FAD → $FADH_2$ |
| **FRD (FrdABCD)**<br>FrdA: P00363 | FAD | Couples menaquinol oxidation to fumarate reduction, producing succinate during anaerobic respiration.<br>Flavin oxidation during fumarate reduction:<br>$FADH_2$ → FAD |
| **Mqo**<br>P33940 | FAD | Oxidizes malate to oxaloacetate using quinone as the electron acceptor.<br>Flavin reduction: FAD → $FADH_2$ |

| **Enzyme or complex** UniProt accession(s) | **Flavin cofactor** | **Metabolic function** |
|---|---|---|
| **NDH-1** NuoF: P31979 | FMN | Transfers electrons from NADH through FMN and iron–sulfur centers to quinones. NuoF is the FMN-binding subunit of the complex. Flavin reduction: FMN → $FMNH_2$ |
| **NDH-2 (Ndh)** P00393 | FAD | Oxidizes NADH and reduces quinones without directly coupling the reaction to proton translocation. Flavin reduction: FAD → $FADH_2$ |
| **Dld** P06149 | FAD | Oxidizes D-lactate to pyruvate and transfers electrons to the quinone pool. Flavin reduction: FAD → $FADH_2$ |
| **LldD** P33232 | FMN | Oxidizes L-lactate to pyruvate and supplies electrons to the respiratory chain. Flavin reduction: FMN → $FMNH_2$ |

[a] Sequence analysis suggests FAD binding by GlpA and FMN binding by GlpB[14].
